# Intercellular compartmentation of trehalose 6-phosphate metabolism in *Setaria viridis* leaves

**DOI:** 10.1101/2024.08.16.608293

**Authors:** Tomás Tonetti, Bruno E. Rojas, Regina Feil, Camila Seimandi, Leandro E. Lucero, Julieta V. Cabello, Paula Calace, Mariana Saigo, Stéphanie Arrivault, Mark Stitt, John E. Lunn, Carlos M. Figueroa

**Author notes:** Equal contribution (Tomás Tonetti, Bruno E. Rojas).

## Abstract

Trehalose 6-phosphate (Tre6P) is a signal metabolite that links carbon metabolism with plant development. Our current understanding of Tre6P metabolism and signalling is predominantly based on studies in *Arabidopsis thaliana*. Some features could be adapted to the specific physiology, anatomy, and life cycle of this C_3_ eudicot model species, and thus might not be representative of other angiosperms. To better understand Tre6P metabolism in monocot C_4_ species we used *Setaria viridis*, which has been widely adopted as a model for the major C_4_ NADP-malic enzyme subtype crop species, such as maize (*Zea mays*), sorghum (*Sorghum bicolor*) and sugarcane (*Saccharum officinarum*). In this work, we analysed the levels of transcripts encoding Tre6P-related enzymes in different tissues and cell types from *S. viridis*. The *TREHALOSE-6-PHOSPHATE SYNTHASE1* transcript, its encoded protein (TPS1, the enzyme responsible for Tre6P synthesis) and Tre6P were mainly located in bundle sheath cells of *S. viridis*. Our results show that Tre6P is predominately synthesized and located in bundle sheath and associated cells.

**Highlight:** Trehalose 6-phosphate, a sugar signalling metabolite, is mainly present in the bundle sheath cells of *Setaria viridis* leaves.

## Introduction

Trehalose is a non-reducing disaccharide widely distributed in nature (Figueroa and Lunn, 2016). In plants, the pathway of trehalose synthesis occurs through the phosphorylated intermediate trehalose 6-phosphate (Tre6P), which is synthesized from UDP-glucose (UDPGlc) and glucose 6-phosphate (Glc6P) by Tre6P synthase (TPS; EC 2.4.1.15) and dephosphorylated to trehalose by Tre6P phosphatase (TPP; EC 3.1.3.12; Cabib and Leloir, 1958; Avonce *et al*., 2006). Trehalose is then hydrolysed to glucose by trehalase (TRE; EC 3.2.1.28). Gene families coding for TPS and TPP are present in species from all of the main taxonomic groups of green plants (Avonce *et al*., 2006; Lunn, 2007; Lunn *et al*., 2014; Figueroa and Lunn, 2016). Plant TPSs are divided into two distinct clades: class I and class II. Class I TPSs are responsible for Tre6P synthesis, while class II TPSs (which lack some of the glycosyltransferase active site residues involved in substrate binding and catalysis) are putatively involved in Tre6P perception and/or signalling (Avonce *et al*., 2006; Lunn, 2007; Vandesteene *et al*., 2010; Lunn *et al*., 2014; Delorge *et al*., 2015; Figueroa and Lunn, 2016; Van Leene *et al.,* 2022). In the last two decades, studies performed with mutant and transgenic lines demonstrated the importance of Tre6P for all stages of plant development (Eastmond *et al*., 2002; van Dijken *et al*., 2004; Gomez *et al*., 2006; Gomez *et al*., 2010; Wahl *et al*., 2013; Fichtner *et al*., 2020; Ponnu *et al*., 2020; reviewed in Fichtner and Lunn, 2021).

In *Arabidopsis thaliana* leaves, Tre6P is synthesized primarily in phloem parenchyma (bundle sheath) cells and in the companion cell-sieve element complex of the phloem (Fichtner *et al*., 2020). The phloem parenchyma cells are symplastically connected with mesophyll cells (MC) via plasmodesmata, allowing Tre6P to diffuse into the MC to regulate sucrose production (Fichtner and Lunn, 2021). Tre6P regulates the synthesis and transport of sucrose by influencing photoassimilate partitioning between sucrose or organic and amino acids during the day (Figueroa *et al*., 2016), transitory starch degradation during the night and towards the end of the light period (Martins *et al*., 2013; dos Anjos *et al*., 2018; Ishihara *et al*., 2022), and expression of *SWEET* (*SUGARS WILL EVENTUALLY BE EXPORTED TRANSPORTER*) genes, which encode transporters involved in sucrose phloem loading (Oszvald *et al*., 2018; Fichtner *et al*., 2021). Therefore, Tre6P coordinates the synthesis and transport of sucrose and amino acids from source leaves with the demand from growing and sink tissues (Martins *et al*., 2013; Figueroa *et al*., 2016; dos Anjos *et al*., 2018). In addition, Tre6P exerts widespread transcriptional repression of photosynthesis, gluconeogenesis and carbon starvation responses, induces amino acid synthesis, nucleotide synthesis and ribosome formation and interacts with light signaling and many phytohormone signaling pathways (Avidan *et al*., 2024).

Our current understanding of Tre6P metabolism and signalling in plants is mainly based on studies in *A. thaliana* (Fichtner and Lunn, 2021; Rojas *et al*. 2023). This model eudicot is a rosette-forming, annual weed species that performs C_3_ photosynthesis and grows predominantly in temperate zones. Some features of Tre6P metabolism and signalling in *A. thaliana* might be adapted to its specific physiology, anatomy and life cycle, and so might differ from other species. Tre6P metabolism and signalling have also been investigated in maize (*Zea mays*), a domesticated monocot species that performs C_4_ photosynthesis and originated in subtropical regions. These studies have revealed the importance of Tre6P metabolism and signalling in the coordination of metabolism in source and sink zones, inflorescence and shoot branching, floral identity, tillering, seed development and tolerance to water deficit (Satoh-Nagasawa *et al*., 2006; Henry *et al*., 2014, 2015; Nuccio *et al*., 2015; Czedik-Eysenberg *et al*., 2016; Bledsoe *et al*., 2017; Oszvald *et al*., 2018; Claeys *et al*., 2019; Dong *et al*., 2019; Klein *et al*., 2022). However, how far Tre6P metabolism and signalling are adapted to the specialized C_4_ photosynthetic metabolism and leaf anatomy of this species is poorly understood.

Tre6P regulates photoassimilate partitioning between sucrose and organic acids and amino acids in *A. thaliana* via coordinated changes in the activities of nitrate reductase, phospho*enol*pyruvate carboxylase (PEPC) and, by inference, the mitochondrial pyruvate dehydrogenase (Figueroa *et al*., 2016). Artificially increased levels of Tre6P reduced the mono-ubiquitination and increased the phosphorylation status of PEPC, thus activating the enzyme. These changes increased carbon flux into organic acids and the TCA cycle to provide carbon skeletons for amino acid synthesis. Given that PEPC is responsible for primary carbon fixation in the C_4_ pathway, Tre6P might therefore play a direct role in regulating photosynthetic carbon fixation in C_4_ plants. This raises the question of whether Tre6P is also involved in the regulation of photoassimilate partitioning to amino acids in C_4_ plants and, if so, how flux into metabolites in the C_4_ cycle and flux into the TCA cycle and amino acid synthesis are reconciled. Tre6P also regulates the mobilization of transitory starch reserves in *A. thaliana* leaves and provision of sucrose at night (Martins *et al*., 2013; dos Anjos *et al*., 2018; Ishihara *et al*., 2022). In contrast to C_3_ plants, where sucrose and starch synthesis both occur within MC, in C_4_ plants starch synthesis is restricted to the bundle sheath cells (BSC), while sucrose is made predominantly in MC (Lunn and Furbank, 1997; 1999). If Tre6P also regulates photoassimilate partitioning and transitory starch turnover in C_4_ plants, this cellular compartmentation of sucrose and starch metabolism would add a further layer of complexity. It has been proposed that expression of the genes encoding the SWEET and SUT/SUC (sucrose-H^+^ symporters) transporters had to be adapted to the specialized Kranz anatomy in C_4_ species, and was potentially a critical element in the evolution of C_4_ plants (Furbank and Kelly, 2021). Mutation of the three maize *ZmSWEET13* genes produced plants with impaired sugar phloem loading and drastic effects on growth (Bezrutczyk *et al*., 2018). As Tre6P has been implicated in the transcriptional regulation of *SWEET* genes (Oszvald *et al*., 2018; Fichtner *et al*., 2021), it is possible that Tre6P also plays a key role in modulating phloem loading during the establishment of Kranz anatomy in developing leaves of C_4_ species.

Currently, we have little knowledge about Tre6P signalling in the leaves of C_4_ plants (Rojas *et al*., 2023). By extrapolation from its known functions in leaves of C_3_ plants, we hypothesize that Tre6P plays a role in the regulation of carbon fixation, photoassimilate partitioning and phloem loading in the leaves of C_4_ plants. If so, Tre6P signalling in these species may have unique features, reflecting their specialized metabolism and anatomy. For instance, the *TREHALOSE-6-PHOSPHATE SYNTHASE1* transcript was mainly found in BSC of several C_4_ species (Benning *et al*., 2023). To better understand how Tre6P signalling operates in C_4_ crops, we investigated the compartmentation of Tre6P in leaves of *Setaria viridis* (wild foxtail millet), a model species for translational research in plants performing NADP-dependent malic enzyme (NADP-ME) C_4_ photosynthesis (Brutnell *et al*., 2010; Doust *et al*., 2019). Our results show that the *TPS1* transcript, its encoded protein (TPS1) and Tre6P are mainly located in BSC.

## Materials and methods

### Plant growth and harvest conditions

Seeds of *Setaria viridis* (L.) P. Beauv. accession A10.1 were obtained from the Germplasm Resources Information Network (GRIN; http://www.ars-grin.gov/), United States Department of Agriculture (PI 669942). To increase the germination rate, the seed coat was abraded with sandpaper (Van Eck and Swartwood, 2015). Seeds were surface sterilized with 0.55 g l^-1^ sodium hypochlorite containing 0.1% (v/v) Tween-20 for 5 minutes, followed by three washes with sterile deionized water. The seeds were then sown on plates with half-strength Murashige and Skoog medium (PhytoTech Labs; https://phytotechlab.com/) supplemented with 0.8% (w/v) agar. Plates were placed in a growth chamber at 28°C with an irradiance of 120 µmol m^-2^ s^-1^, provided by white fluorescent tubes, under a long-day photoperiod regime (16 h light/8 h dark). After five days, seedlings were transferred to individual 8-cm diameter pots containing Klasmann TS1 substrate and irrigated with 1X Hoagland solution. For the analysis of transcripts and proteins, plants were grown in a Conviron Adaptis A1000 controlled environment chamber (https://www.conviron.com/) with an irradiance of 350 µmol m^-2^ s^-1^, provided by white fluorescent tubes, with a long-day photoperiod regime, at 28/22°C (day/night), and 50% relative humidity for 2 weeks. For metabolite fractionation experiments, plants were grown in a walk-in phytotron with an irradiance of 550 µmol m^-2^ s^-1^, provided by LEDs, at 28/22°C (day/night) and 65% relative humidity. Harvested plants had eight fully developed leaves and inflorescences were not macroscopically visible. The blades from the 5^th^ and 6^th^ true leaves (counting from the bottom of the plant) were harvested in the middle of the photoperiod. The leaf material used for isolation of MC and BSC was immediately processed and then stored at −80°C. For enzymatic and metabolite analyses, leaf samples were rapidly quenched in liquid nitrogen and stored at −80°C until use. Samples for the analysis of transcripts and proteins (seedlings, inflorescences, the fourth internode and the flag leaf) were frozen in liquid nitrogen and stored at −80°C until use.

### Phylogenetic analysis

TPS, TPP and TRE protein sequences from *S. viridis* (genome version 2.1) were identified using the keyword “trehalose” at the Phytozome 13 server (https://phytozome.jgi.doe.gov/). We followed the same approach to retrieve TPS and TPP protein sequences from *Ananas comosus* (genome version 3) and *Musa acuminata* (genome version 1). Sequences were manually curated to remove incomplete and duplicated entries. TPS and TPP protein sequences from *A. thaliana*, *Brachypodium distachyon*, *Oryza sativa*, *Phaseolus vulgaris*, *Populus trichocarpa*, *Triticum aestivum* and *Z. mays* were obtained from previous publications (Yang *et al*., 2012; Henry *et al*., 2014; Barraza *et al*., 2016; de Haro *et al*., 2019; Wang *et al*., 2019; Du *et al*., 2022; Griffiths *et al*., 2025). Sequences were aligned with the CLUSTALW tool available at the GenomeNet server (https://www.genome.jp/tools-bin/clustalw). Distance-based phylogenetic trees were reconstructed with the SeaView software version 4.3.0 (Gouy *et al*., 2010) using the BioNJ algorithm (Gascuel, 1997) and 1000 replicates. Figures were prepared with the FigTree program version 1.3.1 (http://tree.bio.ed.ac.uk/software/figtree/). Sequences used to reconstruct the trees were labelled as shown in Supplementary Table S1.

### Separation of mesophyll and bundle sheath cells

Fractions highly enriched in MC or BSC strands were obtained from fresh leaves following the methods described by John *et al*. (2014), with the modifications introduced by Calace *et al*. (2021). In both cases, each replicate consisted of the material obtained from 20 plants. These samples were used for the analysis of transcripts and proteins. To assess the suitability of the fractionation procedure, we performed western blots using commercial antibodies raised against the large subunit of Rubisco (RbcL; Agrisera AS03 037; https://www.agrisera.com/) and PEPC (Agrisera AS09 458), as described below for the immunoblotting of SvTPS1. Supplementary Fig. S1 shows that Rubisco and PEPC were exclusively detected in BSC and MC fractions, respectively.

To determine the distribution of metabolites, frozen leaf tissue was finely ground in liquid nitrogen and the frozen tissue powder was passed through a series of nylon meshes of different pore sizes (200, 80, and 41 µm). The material that was retained on each filter and the one that passed through the 41 µm filter were collected to obtain the samples named F200, F80, F40, and FT, respectively. The F200 fraction, depending on the grinding, often has larger whole leaf bits but tends to be enriched in BSC; the F80 fraction is especially enriched in BSC; the F40 fraction is usually a mix of BSC and MC; and FT fraction is especially enriched in MC (Stitt and Heldt, 1985). This method is suitable for the quantification of metabolites with short half-lives (Arrivault *et al*., 2009, 2017). Metabolites and enzymes, including cell-specific marker enzymes, were measured in each fraction, and their distribution between MC and BSC were calculated by linear regression, as described by Stitt and Heldt (1985). Two C_4_ photosynthetic enzymes were used as markers for each cell-type: PEPC and NADP-dependent malate dehydrogenase (NADP-MDH) for MC and NADP-ME and phosphoribulokinase (PRK) for BSC. Each biological replicate consisted of the material obtained from 20 plants.

### Total RNA extraction and cDNA synthesis

Total RNA was extracted from 50 mg fresh weight (FW) of finely ground tissue with the TransZol reagent (TransGen Biotech; https://www.transgenbiotech.com/), following the manufacturer’s instructions. After RNA extraction, samples were treated with DNase (Promega; https://www.promega.com) and RNA concentration was quantified using a NanoDrop 2000 (Thermo Scientific; https://www.thermofisher.com). cDNA was synthesized with the MultiScribe Reverse Transcriptase (Applied Biosystems; https://www.thermofisher.com), according to the manufacturer’s instructions.

### Determination of transcript levels

Real-time reverse transcriptase-qPCR (RT-qPCR) reactions were performed in a final volume of 8 µl in 96-well microplates (Applied Biosystems) with the iQ SYBR Green Supermix (Bio-Rad; https://www.bio-rad.com), 0.2 µM of each primer, and an appropriate dilution of the sample. Transcript abundance was calculated with the comparative C_t_ method (Schmittgen and Livak, 2008) and normalization was performed with the *SvKIN* transcript (Sevir.4G252700; Martins *et al*., 2016), which showed similar C_t_ values across all the analysed samples (Supplementary Fig. S2). All the primers used in this work are listed in Supplementary Table S2.

### Production of a polyclonal antiserum against SvTPS1

The protein coding region of the class I *TPS* gene (SvTPS1; Uniprot ID A0A4U6VAG9) was codon-optimised and *de novo* synthesized (BioBasic; https://www.biobasic.com/) for expression in *E. coli* (Supplementary File S1) and cloned into the pET28b expression vector (Novagen; https://www.merckmillipore.com) between the NdeI and SacI restriction sites, in frame with a His_6_-tag at the N-terminus. The resulting vector was used to transform *E. coli* BL21 (DE3) cells (Invitrogen; https://www.thermofisher.com). Transformed cells were grown in Lysogeny Broth medium, supplemented with 50 µg ml^-1^ kanamycin at 37°C, with shaking at 200 rpm until the optical density at 600 nm reached approximately 0.6. Synthesis of the recombinant protein was induced by addition of isopropyl-β-D-thiogalactopiranoside to a final concentration of 1 mM and cells were incubated for 16 h at 25°C and 200 rpm after induction. The recombinant SvTPS1 protein was mainly present in the insoluble fraction; thus, it was purified from inclusion bodies (Jenö and Horst, 2002). This protein was used to raise a polyclonal antiserum in rabbits at ICIVET Litoral (UNL-CONICET, Argentina). Antibodies were affinity-purified using a nitrocellulose membrane strip containing the recombinant SvTPS1, as described by Fang (2012).

### Protein extraction

Plant material was homogenized in a pre-cooled mortar with liquid nitrogen. For denaturing protein extraction, 20 mg FW of frozen tissue were extracted with 200 µl of 125 mM Tris-HCl pH 6.8, 2% (w/v) sodium dodecyl sulphate (SDS), 20% (v/v) glycerol, 1.4 M 2-mercaptoethanol, and 0.05% (w/v) Bromophenol Blue. After adding the buffer, samples were vortex-mixed and heated for 5 min at 95°C with agitation. Samples were then cooled to room temperature and centrifuged at 20,000 x *g* for 10 min to separate the protein extract from insoluble tissue debris. Alternatively, 50 mg FW of frozen tissue were resuspended in 500 µl of 10 mM Tris-HCl pH 8.0, 10% (v/v) trichloroacetic acid (TCA), 25 mM ammonium acetate, and 1 mM EDTA (Rojas *et al*., 2021). Samples were incubated at 4°C for 30 min and centrifuged at 20,000 x *g* for 15 min. The supernatant was discarded and the pellet was washed with 500 µl of 5% (w/v) TCA and then with 500 µl of 90% (v/v) acetone, then allowed to dry before the protein pellet was dissolved in 50 µL of Laemmli buffer for analysis by SDS-PAGE.

Protein extraction for enzymatic assays was performed as described by Gibon *et al*. (2004), with minor modifications. Samples (20 mg FW of frozen tissue) were supplemented with 10 mg of polyvinylpolypyrrolidone and then resuspended in 500 µl of 50 mM HEPES-KOH pH 7.5, 10 mM MgCl_2_, 1 mM EDTA, 1 mM EGTA, 1 mM benzamidine, 1 mM ε-aminocaproic acid, 0.02 mM leupeptin, 1 mM DTT, 1 mM phenylmethylsulfonyl fluoride, 0.1% (v/v) Triton X-100, and 20% (v/v) glycerol. Samples were incubated on ice for 10 min and then centrifuged at 20,000 x *g* and 4°C for 10 min to separate soluble proteins from tissue debris. Protein concentration was determined with the Bradford reagent (Bradford, 1976), using a standard curve constructed with bovine serum albumin.

### Immunoblotting

Protein extracts were subjected to a 10% SDS-PAGE and electrotransferred to a 0.45 µm nitrocellulose membrane (Amersham; https://www.cytivalifesciences.com/). To check lane loadings, membranes were stained with 0.01% (w/v) Ponceau S in 1% (v/v) acetic acid. After washing with TBST1 solution [50 mM Tris-HCl pH 8.0; 150 mM NaCl; 0.2% (v/v) Tween-20], membranes were blocked for 1 h with TBST1 supplemented with 0.2% (w/v) fat-free dried milk powder. Membranes were incubated overnight at 4°C with purified anti-SvTPS1 in TBST1 supplemented with 0.2% (w/v) fat-free dried milk powder. After washing, the membranes were incubated for 30 min at room temperature with goat anti-rabbit IgG (H&L) conjugated to horseradish peroxidase (Agrisera AS09 602). Proteins were detected using the ECL Bright reagent (Agrisera AS16 ECL-N) and exposed to X-ray films (AGFA; https://www.agfa.com/corporate/) for 1 min. The sizes of the immunoreactive bands were estimated from the relative mobility (Rf) *versus* logarithm of molecular weight (logMM) plot constructed with protein standards, using data from three independent experiments.

### Immunoprecipitation of SvTPS1

Proteins were extracted from 100 mg FW of BSC with 400 μl of a buffer containing 100 mM Bicine pH 9.0, 10% (v/v) glycerol, 0.1% (v/v) Triton X-100, 5 mM 2-mercaptoethanol, 1 mM EDTA, 1 mM EGTA, 2 mM phenylmethylsulfonyl fluoride and 1 X Protease Inhibitor Cocktail III (Calbiochem; https://www.sigmaaldrich.com/). Immunoprecipitation was performed as described by Hartman *et al*. (2023), with minor modifications. Briefly, antibody-bead conjugates were prepared by adding 5 μl of polyclonal anti-SvTPS1 antibody to a microcentrifuge tube containing 200 µl of TBST2 [50 mM Tris-HCl pH 8.0; 150 mM NaCl; 0.05% (v/v) Tween-20] and 75 μl of protein A-Sepharose beads (Sigma). Conjugates were incubated 90 min at room temperature with gently mixing and then washed with TBST2 to remove unbound antibodies. The protein extract (150 μl) was incubated with the antibody-bead conjugate for 16 h at 4°C with gently mixing. After washing with TBST2, the pellet was resuspended with Laemmli buffer and heated at 95°C for 10 min. The sample was then centrifuged at room temperature and 10,000 x *g* for 5 min and the resulting supernatant was separated by SDS-PAGE. The gel was stained using colloidal Coomassie Blue G-250. Controls were processed in parallel either without antibodies or the protein extract.

### Mass spectrometry analysis

Tryptic peptides from the gel fragment were obtained using the protocol of Link and LaBaer (2009) and analysed by mass spectrometry (MS) at the Mass Spectrometry Unit of the Institute of Molecular and Cellular Biology of Rosario (UNR-CONICET), Argentina. The sample (3 µl) was injected on an nanoHPLC Ultimate3000 (Thermo Scientific) and peptides were separated using a nano column EASY-Spray ES901 (15 cm × 50 μm ID, PepMap RSLC C18). The mobile phase flow rate was 300 nl/min using 0.1% formic acid in water (solvent A) and 0.1% formic acid and 100% acetonitrile (solvent B). The gradient profile was set as follows: 4-30% solvent B for 10 min, 30%-80% solvent B for 2 min and 90% solvent B for 1 min. MS analysis was performed using a Q-Exactive HF mass spectrometer (Thermo Scientific). For ionization, 1,9 kV of liquid junction voltage and 250°C of capillary temperature were used. The full scan method employed a m/z 375–2000 mass selection, an Orbitrap resolution of 120,000 (at m/z 200), a target automatic gain control (AGC) value of 1e6 and a maximum injection time of 100 ms. After the survey scan, the 5 most intense precursor ions were selected for MS/MS fragmentation.

Fragmentation was performed with a normalized collision energy of 27 eV and MS/MS scans were acquired with a dynamic first mass, AGC target was 5e5, resolution of 30000 (at m/z 200), isolation window of 1.4 m/z units and maximum IT was 55 ms. Charge state screening was enabled to reject unassigned, singly charged, and equal or more than six protonated ions. A dynamic exclusion time of 10 s was used to discriminate against previously selected ions.

MS data were analysed with Proteome Discoverer 2.4.1.15, using standardized workflows. The mass spectra raw file was searched against the *Setaria viridis* deduced proteome (UniProt ID UP000298652). Precursor and fragment mass tolerance were set to 10 ppm and 0.02 Da, respectively, allowing 2 missed cleavages, with Cys carbamidomethylation (+57.021 Da) as fixed modification, and the following dynamic modifications: Met oxidation (+15.995 Da), N-term acetylation (+42.011 Da), Met loss (−131.040 Da) and Met loss plus acetylation (−89.030 Da). The raw proteomics data were deposited to the PRIDE repository (https://www.ebi.ac.uk/pride/) with the dataset identifier PXD057139.

### Immunohistochemistry

Tissue fixation, dehydration, assembly of the paraffin block and microtome cutting were performed as described by Cabello and Chan (2019), with minor modifications. Sections of 0.5-1.0 cm in length from the sixth leaf of three-week-old plants (without the midrib) were fixed at 25°C for 1 h in a solution containing 3.7% (v/v) formaldehyde, 5% (v/v) acetic acid and 47.5% (v/v) ethanol. To further fix and discolour the tissue, sections were incubated in 70% (v/v) ethanol for two weeks. Sections were dehydrated through a series of ethanol solutions (80, 90, 96 and 100%; 30 min each), followed by incubation in ethanol-xylene solutions (3:1; 1:1 and 1:3; 1 h each), and finally stored overnight in 100% xylene at room temperature. To obtain the paraffin blocks, sections were incubated through a series of xylene-paraffin solutions (3:1, 1:1 and 1:3; 1 h each). The samples were placed into plastic casts and finally embedded in Histoplast (Biopack; https://www.biopack.com.ar/). Each block was incubated overnight at room temperature to ensure solidification. Cross-sections (10 μm thick) were obtained using a Microtome RM2125 (Leica Microsystems; https://www.leica-microsystems.com/) and mounted on slides coated with 50 mg/ml poly-D-Lys (Sigma) in 10 mM Tris-HCl pH 8.0 and dried for 16 h at 37°C. Paraffin was removed with 100% xylene for 15 min at room temperature and sections were rehydrated using a series of ethanol (100, 96, 90, 80, 70 and 50%; 1 min each) to finish in distilled water.

Immunolocalization of proteins in *S. viridis* leaves was performed as described by Sauer and Friml (2010), with minor modifications. Mounted cross-sections were rinsed three times in PBS (10 mM sodium phosphate, 130 mM NaCl, pH 7.4) and blocked with PBS with the addition of 1% (w/v) BSA and 0.1% (v/v) Tween-20 (blocking solution) at room temperature for 1 h in a humid chamber. The blocking solution was removed and sections were incubated for 2 h at room temperature with the anti-RbcL (Agrisera AS03 037; dilution 1:1000) or the purified anti-SvTPS1 (see above; dilution 1:100) in blocking solution. Tissue sections were washed three times with blocking solution and then incubated with Cy2-conjugated goat anti-rabbit IgG (Jackson ImmunoResearch 111-225-144; https://www.jacksonimmuno.com/; dilution 1:200). Samples were incubated at room temperature for 1 h, washed three times with blocking solution and air-dried. Sections incubated only with the secondary antibody were used as a negative control. Finally, samples were mounted with 85% (v/v) glycerol and a coverslip. Micrographs were taken with a SP8 LIGHTNING confocal microscope (Leica Microsystems) using the 20X objective and the Cy2 bis NHS ester fluorophore channel (excitation 492 nm, emission 510 nm). Images were processed using the LAS X Office 1.4.7.28982 software (Leica Microsystems) and figures were prepared with Inkscape 0.92.2 (https://inkscape.org/).

### Metabolite extraction and measurement

Phosphorylated intermediates and organic acids were extracted with a mixture of chloroform:methanol (3:7) and then measured by anion-exchange high-performance liquid chromatography coupled to tandem mass spectrometry (LC-MS/MS), as described by Lunn *et al*. (2006), with the modifications introduced by Figueroa *et al*. (2016). Calvin-Benson cycle intermediates and aspartate were measured in chloroform-methanol extracts by ion-pair reverse phase LC-MS/MS, as described by Arrivault *et al*. (2009). Pyruvate, phospho*enol*pyruvate (PEP), dihydroxyacetone phosphate (DHAP) and 3-phosphoglycerate (3PGA) were enzymatically measured in TCA extracts (Arrivault *et al*., 2017). The content of sucrose, glucose, and fructose in ethanolic extracts was determined following the method of Stitt *et al*. (1989), while the amount of starch was measured using the insoluble residue from the ethanolic extraction, as described by Hendriks *et al*. (2003). The datasets containing metabolite levels and the associated calculations are available in Supplementary Tables S3-S5.

### Enzyme activity assays

PRK activity was measured spectrophotometrically using a continuous assay (Leegood, 1990), with minor modifications. The assay mixture contained: 50 mM Tricine-KOH pH 8.0, 1 mM ATP, 50 mM KCl, 0.3 mM NADH, 10 mM MgCl_2_, 2 mM PEP, 2 U pyruvate kinase, and 2 U lactate dehydrogenase and the reaction was started by adding 0.5 mM ribulose 5-phosphate. PEPC, NADP-MDH, and NADP-ME activities were measured as described by Ashton *et al*. (1990), with minor modifications. The PEPC assay mixture contained: 50 mM Tricine-KOH pH 8.0, 5 mM MgCl_2_, 2 mM DTT, 1 mM KHCO_3_, 5 mM Glc6P, 0.3 mM NADH, and 0.4 U malate dehydrogenase and the reaction was started with 2 mM PEP. The NADP-MDH assay mixture contained: 50 mM Tricine-KOH pH 8.0, 70 mM KCl, 1 mM EDTA, 1 mM DTT, and 0.3 mM NADPH and the reaction was started with 1 mM oxaloacetate (freshly prepared). The NADP-ME assay mixture contained: 50 mM Tricine-KOH pH 8.0, 0.5 mM NADP^+^, 0.1 mM EDTA, and 2 mM MgCl_2_ and the reaction was started with 5 mM malate. All reactions were performed in a final volume of 200 µl at 25°C in 96-well microplates. The reduction/oxidation of NAD(P)^+^/NAD(P)H was followed by measuring the absorbance at 340 nm in a Biotek ELx808 microplate reader. The amount of crude extract added to the reaction was chosen to ensure a linear response of the signal. One unit (U) of enzyme activity is defined as the amount of extract producing 1 µmol of product in 1 min under the specified assay conditions.

## Results

### Phylogenetic analysis of Tre6P-related enzymes

To identify the proteins involved in Tre6P metabolism in *S. viridis*, we performed keyword searches against its deduced proteome. We found 21 putative Tre6P-related enzymes: 10 TPSs, 10 TPPs, and 1 TRE (Supplementary Table S1), as previously observed for *Setaria italica* (Henry et al., 2020), a domesticated relative of *S. viridis*. After removing duplicated sequences or splicing variants (for SvTPP1, SvTPP8, and SvTPP10), phylogenetic trees were reconstructed for TPS and TPP proteins (Fig. 1 and Supplementary Fig. S3, respectively).

**Figure 1.**
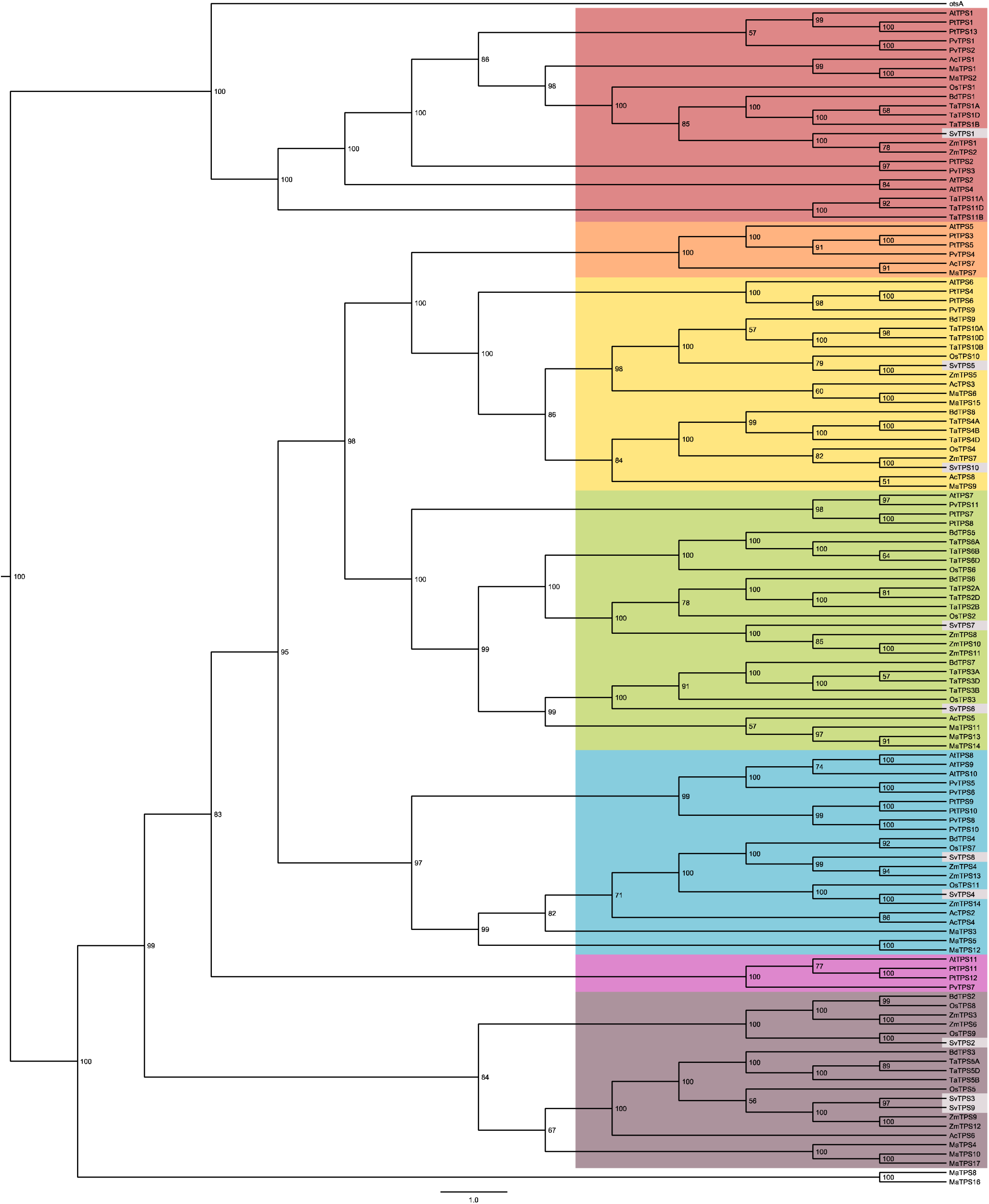
Phylogenetic relationships of *Setaria viridis* TPS proteins. TPS protein sequences were collated from *Setaria viridis* (Sv) and other selected species: Ac, *Ananas comosus*; At, *Arabidopsis thaliana*; Bd, *Brachypodium dystachion*; Mc, *Musa acuminata*; Os, *Oryza sativa*; Pt, *Populus trichocarpa*; Pv, *Phaseolus vulgaris*; Ta, *Triticum aestivum*; Zm, *Zea mays* (see Supplementary Table S1 for gene/protein identifiers). A distance-based tree was reconstructed using the BioNJ algorithm. Numbers in nodes represent the bootstrap obtained for 1000 replicates. The tree is rooted on the *Escherichia coli* TPS (otsA). Sub-clades in the tree are coloured according to the position of *A. thaliana* TPS sequences: AtTPS1/2/4 (red), AtTPS5 (orange), AtTPS6 (yellow), AtTPS7 (green), AtTPS8/9/10 (blue) and AtTPS11 (violet), with the exception of the monocot-specific sub-clade (brown). *S. viridis* sequences are highlighted in grey.

*S. viridis* TPS sequences were split into two major clades containing class I and class II TPSs, as previously described for TPSs from *A. thaliana*, poplar, rice, pea, wheat and maize (Leyman *et al*., 2001; Avonce *et al*., 2006; Lunn, 2007; Yang *et al*., 2012; Henry *et al*., 2014; Barraza *et al*., 2016; Paul *et al*., 2018; de Haro *et al*., 2019). *S. viridis* has only one class I TPS (SvTPS1), which diverges from the other nine class II TPS proteins (SvTPS2-10; Fig. 1). All *S. viridis* TPS sequences displayed the glycosyl-transferase family 20 domain (InterPro ID IPR001830) and the TPP domain (InterPro ID IPR003337) that are typical of plant TPS proteins. Inspection of the primary sequence of *S. viridis* TPSs in detail showed that only SvTPS1 had the characteristic residues involved in UDPGlc and Glc6P binding (Supplementary Fig. S4). The class II TPS sequences from *S. viridis* (SvTPS2-10) were distributed into several branches, most of which contained proteins from eudicot and monocot species. The exceptions were the monocot-specific sub-clade containing SvTPS2/3/9 and that containing *M. acuminata* TPS8 and TPS16, which branched at the base of the class II TPS tree (Fig. 1), an observation initially made by Lunn (2007). A detailed inspection of *S. viridis* class II TPS sequences showed substitutions in the substrate-binding residues of the glucosyltransferase domain that might explain their apparent lack of catalytic activity (Supplementary Fig. S4; Gibson *et al*., 2002; Vandesteene *et al*., 2010). The phylogenetic and sequence analyses indicate that SvTPS1 is the enzyme solely responsible for Tre6P synthesis in *S. viridis*.

The *S. viridis* TPP family was separated into two major clades: three members in clade A and the other seven in clade B (Supplementary Fig. S3). All TPP sequences from *S. viridis* had the TPP domain (InterPro ID IPR003337) and the three conserved active site motifs previously described for these proteins (Avonce *et al*., 2006; Lunn, 2007; Vandesteene *et al*., 2012; Henry *et al*., 2014; Kerbler *et al*., 2023); thus, all 10 *S. viridis* TPP proteins are likely to be catalytically active (Supplementary Fig. S5). Among these, we found SvTPP10 and SvTPP2, the orthologues of RAMOSA3 (ZmTPP10) and ZmTPP4, respectively (Supplementary Fig. S3), which modify inflorescence branching, sex determination and tillering in maize (Satoh-Nagasawa et al., 2006; Claeys et al. 2019; Klein et al. 2022), independently of their catalytic activity.

### Differential expression and compartmentation of Tre6P-related transcripts

To analyse the expression pattern of the identified *S. viridis TPS*, *TPP* and *TRE* genes, we performed real-time RT-qPCR analysis of mRNA extracted from the aerial part of five-days old seedlings, whole leaves (pool of leaves 5 and 6), the flag leaf, whole inflorescences, and the fourth internode of the stem (all samples harvested in the middle of the day). In addition, we measured the levels of transcripts encoding enzymes involved in the C_4_ cycle with differential expression in MC (*SvNADP-MDH* and *SvPEPC*) and BSC (*SvNADP-ME* and *SvRbcS*, which codes for the small subunit of Rubisco) from *S. viridis* and maize leaves (Chang *et al*., 2012; John *et al*., 2014; Li *et al*., 2010). We also analysed the transcripts encoding the sucrose transporters *SvSWEET13a* and *SvSWEET13b*, which have been detected in BSC from *S. viridis* leaves (Chen *et al*., 2022) and are putatively regulated by Tre6P (Oszvald *et al*., 2018; Fichtner *et al*., 2021). These transcripts were detected in all samples, with the highest expression in fully developed leaves and seedlings (Supplementary Fig. S6).

*SvTPS1* was mainly detected in whole leaves and the internode and, to a lesser extent, in seedlings, the flag leaf and the inflorescence. Class II *TPS* transcripts were differentially expressed among the analysed tissues; most of these transcripts were highly expressed in whole leaves and seedlings but only marginally in the internode and the inflorescence. The *SvTPS3* transcript showed the highest level in seedlings and whole leaves; the levels of *SvTPS6* and *SvTPS9* were similar to those observed for *SvTPS1* in whole leaves; the *SvTPS7* transcript displayed the highest level in the flag leaf; whereas the levels of *SvTPS7* and *SvTPS6* were comparable but lower than for *SvTPS1* in the internode (Supplementary Fig. S6).

*SvTPP* transcripts also showed a differential spatial distribution, being higher in whole leaves and internode samples. Among clade A *TPP* transcripts, *SvTPP1* showed the highest levels in whole leaves, the inflorescence and the internode, with a maximum in the latter, while *SvTPP8* displayed the highest levels in seedlings and the flag leaf. Regarding clade B *TPP* transcripts, *SvTPP2* was consistently the one with the highest level in all the samples, showing a maximum in the fourth internode. The highest level of the *SvTPP10* transcript (the orthologue of *RAMOSA3*) was observed in the fourth internode. *SvTRE* was detected in all the samples and its abundance was maximal in the internode (Supplementary Fig. S6).

To determine the compartmentation of these transcripts in mature leaves, we applied fractionation protocols (John *et al*., 2014; Calace *et al*., 2021) to enrich samples in BSC and MC, followed by RNA extraction and analysis by RT-qPCR (Fig. 2). The marker transcripts *SvNADP-MDH* and *SvPEPC* were highly enriched in the MC fraction, while *SvNADP-ME* and *SvRbcS* were mainly detected in the BSC fraction (Fig. 2A). The *SvTPS1* transcript was almost exclusively found in the BSC fraction (Fig. 2B). It should be noted that BSC strands, while enriched in BSC, also contain phloem (companion cells and sieve elements) and xylem tissue. In *A. thaliana* leaves, the AtTPS1 protein is localized primarily in phloem parenchyma (bundle sheath) cells and in the companion cell-sieve element complex of the phloem (Fichtner *et al*., 2020). Thus, some of the *SvTPS1* transcript we attribute to the BSC might be in the phloem complex. The *SvTPS3* and *SvTPS10* transcripts were also preferentially located in the BSC fraction, while *SvTPS6* and *SvTPS8* also showed higher levels in BSC than MC, although with a lower enrichment (Fig. 2B). Conversely, *SvTPS4* and *SvTPS7* transcripts were more abundant in the MC fraction; indeed, *SvTPS7* was not even detected in BSC (Fig. 2B). Among the *TPP* transcripts, *SvTPP1* displayed similar levels in both fractions; *SvTPP2*, *SvTPP3*, *SvTPP7* and *SvTPP9* showed a significant enrichment in the BSC fraction; and *SvTPP4* was only detected in the BSC fraction (Fig. 2C). The *SvTRE* transcript was equally distributed in both cell types, whereas the *SvSWEET13a* and *SvSWEET13b* transcripts were highly enriched in BSC (Fig. 2D). It is important to note that Fig. 2 does not include transcripts that were neither detected in BSC nor MC fractions.

**Figure 2.**
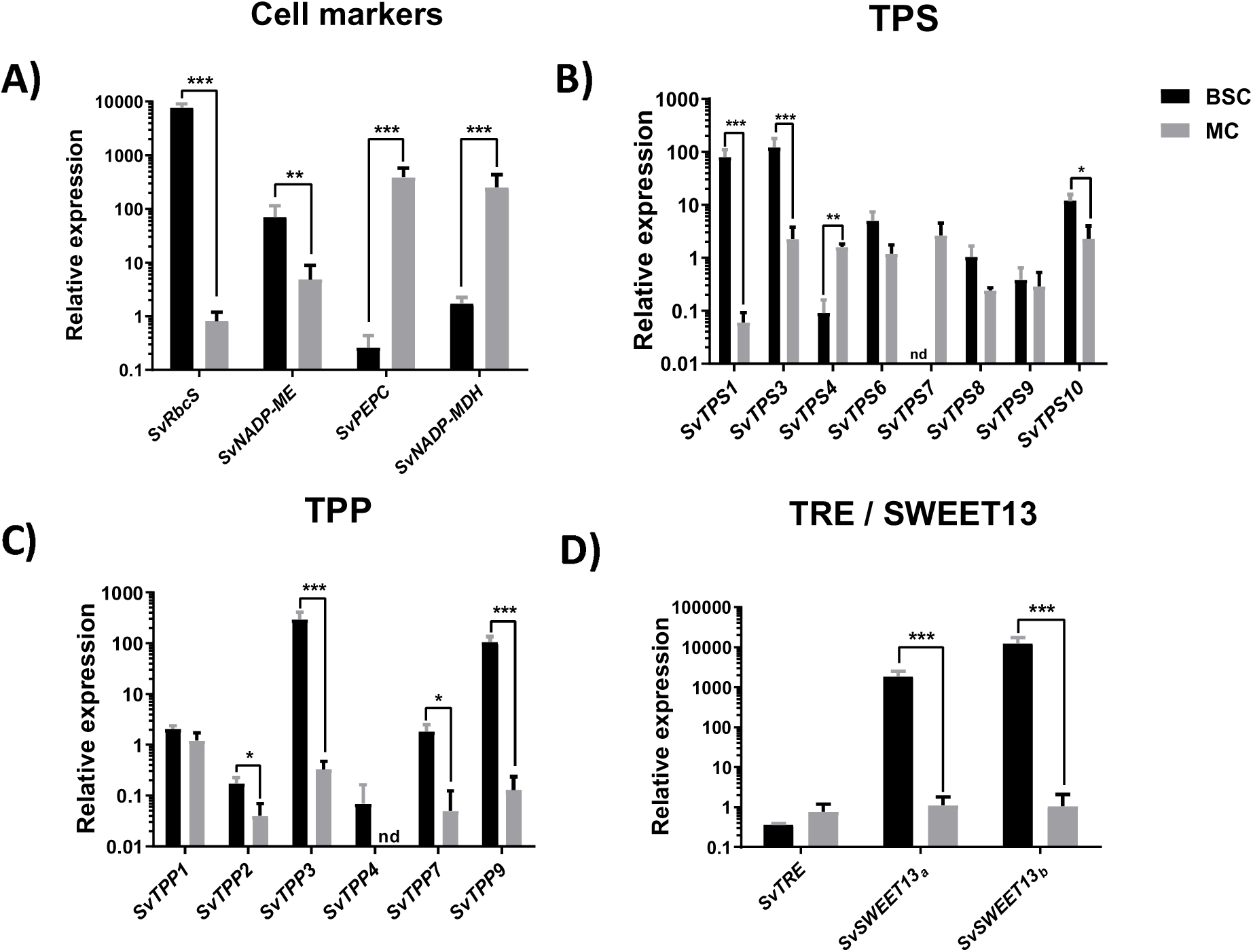
Relative expression of transcripts encoding Tre6P-related enzymes in MC and BSC from *S. viridis* leaves. Relative expression data were calculated with the 2^-ΔCt^ method, using the *SvKIN* transcript as a reference. Data are the mean ± standard error of 4 biological replicates. Statistical analysis was performed using a two-tailed t-test (unequal variance). Significant differences are indicated with asterisks: *, p<0.05; **, p<0.01; ***, p<0.001. nd, not detected.

### Tissue- and cell-specific distribution of the SvTPS1 protein

We recently performed a proteomic analysis with samples enriched in MC and BSC from *S. viridis* leaves (Calace *et al*., 2021). In our dataset (Supplementary Table S1 in Calace *et al*., 2021), which contains 1334 proteins from MC and 1268 proteins from BSC, we observed that SvTPS1 was detected in two out of four biological replicates from the BSC fraction and was absent from the MC fraction (Fig. 3A). To further investigate the distribution of the SvTPS1 protein in different cells, we used an antiserum raised against recombinant SvTPS1 and proteins extracted from MC and BSC enriched samples with TCA and Laemmli buffers. The antiserum recognized two protein bands in the BSC fraction of approximately 112 and 118 kDa (Fig. 3B-C), in agreement with the predicted molecular mass of SvTPS1 (108.4 kDa). Extraction with TCA is expected to rapidly denature any proteases in the samples (Rojas *et al*., 2021), indicating that the double band does not result from proteolytic cleavage of the SvTPS1 protein during the extraction procedure. We then analysed the distribution of SvTPS1 in different tissues. The protein was detected in all the analysed samples, with maximal signals in whole leaves and the internode (Fig. 3D), which resembles the distribution observed for the *SvTPS1* transcript (Supplementary Fig. S6). The loading controls and full-size images that support Figs 3B-D are presented in Supplementary Fig. S7, which also shows that the purified anti-TPS1 antiserum cross-reacted with a protein of ∼50 kDa in samples extracted with Laemmli buffer.

**Figure 3.**
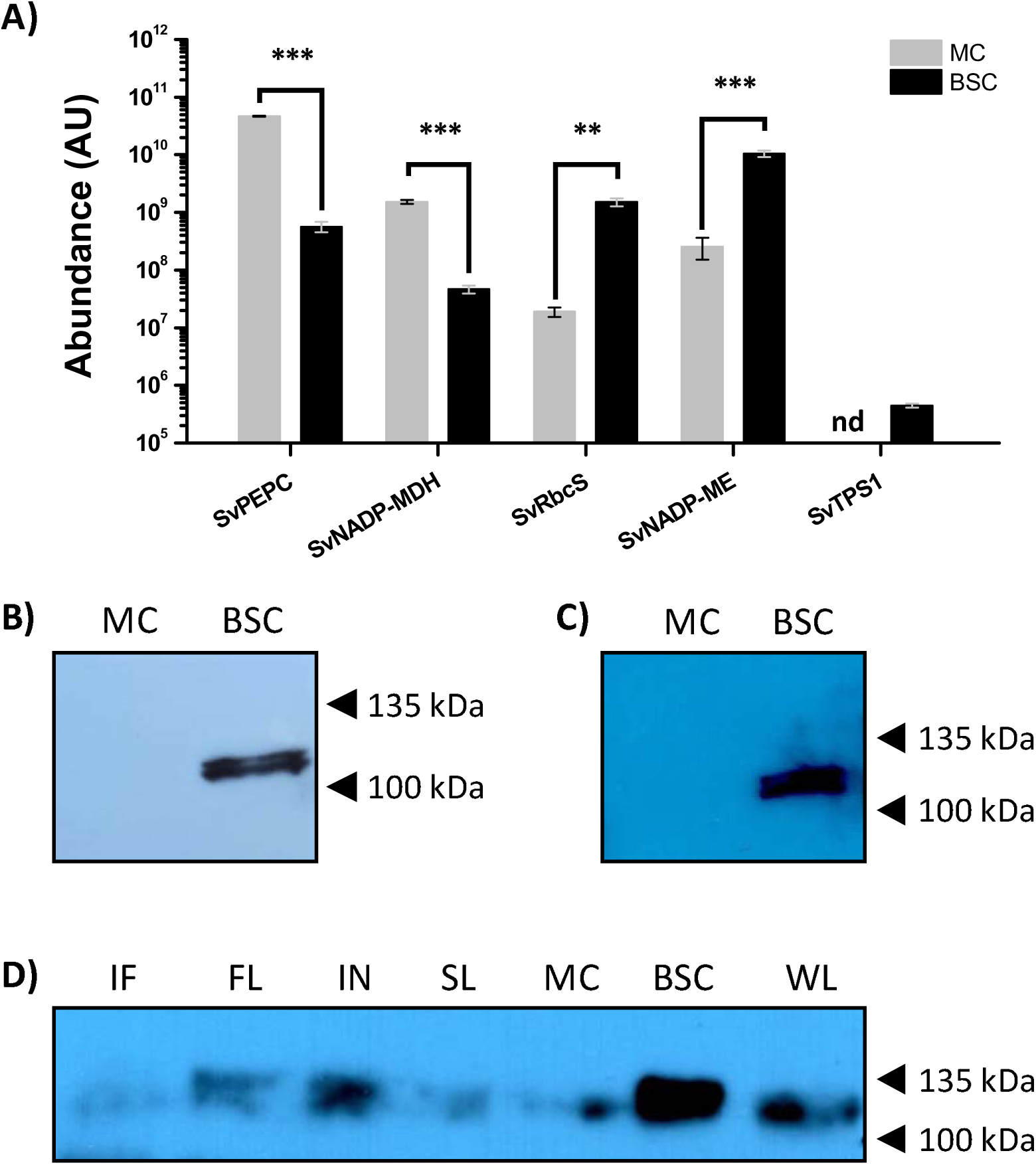
Immunoblotting of SvTPS1 in different tissues and cell types of *S. viridis*. (A) Protein abundance in MC and BSC from *S. viridis* leaves. Data were retrieved from Calace *et. al*. (2021). MC, grey bars; BSC, black bars; AU, arbitrary units; nd, not detected. SvPEPC, Sevir.4G143500; SvNADP-MDH, Sevir.6G256400; SvRbcS, Sevir.3G276100; SvNADP-ME; Sevir.5G132500; SvTPS1, Sevir.3G180900. Data are the mean ± standard error of 4 biological replicates (n=4), except for SvTPS1, which was only detected in two BSC samples (n=2). Statistical analysis was performed using a two-tailed t-test (unequal variance). Significant differences are indicated with asterisks: **, p<0.01; ***, p<0.001. (B) Immunoblotting of SvTPS1 in MC and BSC from *S. viridis* leaves extracted with TCA. Each lane was loaded with extract from 2 mg fresh weight of tissue. (C) Immunoblotting of SvTPS1 in MC and BSC from *S. viridis* leaves extracted with Laemmli buffer. Each lane was loaded with extract from 2 mg fresh weight of tissue. (D) Immunoblotting of SvTPS1 in different tissues and cell types from *S. viridis*. Each lane was loaded with 25 µg of protein. IF; inflorescence, FL; flag leaf, IN; internode, SL; seedlings, MC; mesophyll cells, BSC; bundle sheath cells, WL; whole leaf. The loading controls and full-size images are presented in Supplementary Fig. S7.

To confirm the identity of the protein recognized by the polyclonal antibody, we immunoprecipitated SvTPS1 from samples enriched in BSC. A protein band corresponding to ∼115 kDa was sliced from the gel and analysed by LC-MS (Supplementary Fig. S8). Supplementary Table S6 shows that the most abundant protein from *S. viridis* present in the sample was TPS1 (UniProt ID A0A4U6VAG9), with a coverage of 36%, 28 unique peptides and 51 spectra. We analysed in detail the MS data for immunoprecipitated SvTPS1 but we did not detect any post-translationally modified peptide that could explain the existence of two immunoreactive bands, probably due to the extraction and/or immunoprecipitation procedures employed in this work.

To confirm the location of SvTPS1 on *S. viridis* leaves we performed an immunohistochemical assay. Cross-sections of *S. viridis* leaves were incubated with purified anti-SvTPS1, anti-RbcL (positive control for BSC) or without primary antibody (negative control), and then with goat anti-rabbit IgG conjugated with Cy2. As shown in Fig. 4 and Supplementary Fig. S9, SvTPS1 was preferentially detected in BSC, although we also observed fluorescence in some MC. This result was consistent with the signal observed for the large subunit of Rubisco, which was mainly detected in BSC and, to a lesser extent, in MC (Fig. 4 and Supplementary Fig. S9). The negative control did not show any differential signal (Fig. 4 and Supplementary Fig. S9). Overall, these results indicated that the TPS1 protein is mainly located in BSC of *S. viridis* leaves.

**Figure 4.**
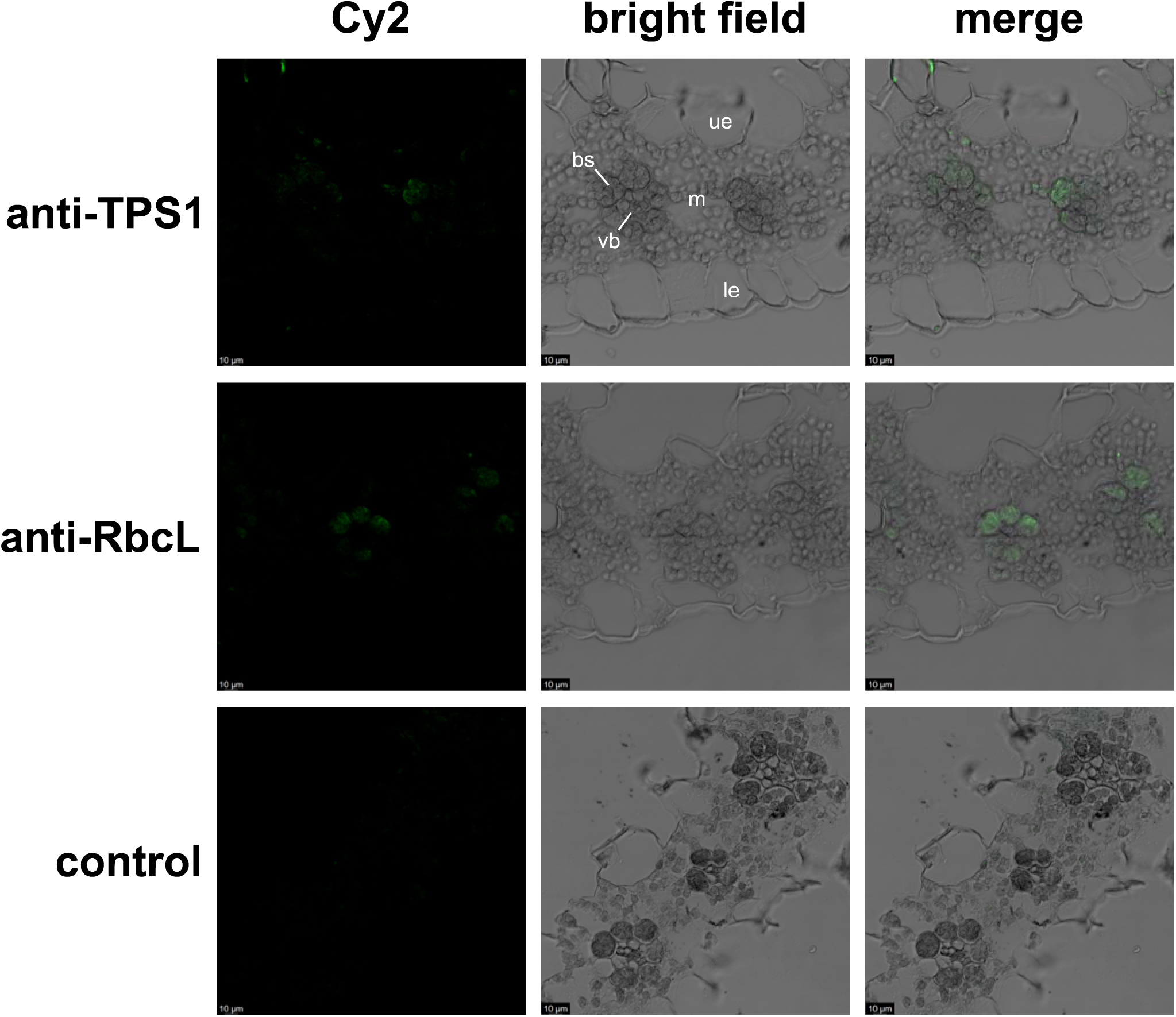
Immunolocalization of SvTPS1 in cross-sections of *S. viridis* leaves. Top row, anti-SvTPS1 (1/100); middle row, anti-RbcL (1/1000); bottom row, negative control (without primary antibody). The secondary antibody (anti-rabbit IgG, 1/200) was conjugated with Cy2. The green channel shows the Cy2 fluorescence and the merged images show the Cy2 signal superimposed on the bright-field images. The panel was prepared with digitally-maximized images (2.5X). Bars=10 µm. Non-maximized images are presented in Supplementary Figure S9. The panel shows representative images of three independent experiments (n=3). bs, bundle sheath; vb, vascular bundle; m, mesophyll; ue, upper epidermis; le, lower epidermis.

### Intercellular distribution of Tre6P

To determine the intercellular distribution of Tre6P, we prepared fractions enriched in MC or BSC from *S. viridis* leaves by sequential filtering of frozen leaf tissue powder through a series of nylon meshes with different pore sizes according to Stitt and Heldt (1985). We performed two independent experiments (Experiments 1 and 2, Supplementary Tables S3 and S4, respectively), each one consisting of four biological replicates. Supplementary Fig. S10 shows the results obtained for marker enzymes from Experiment 1. The recovery of marker enzymes in Experiments 1 and 2 varied between 57 and 87%, depending on the enzyme and the experiment (Supplementary Table S7). The relative distribution of these enzymatic activities indicated that the fractionation method was successfully adapted to *S. viridis* leaves, as the results resemble those obtained for maize leaves (Stitt and Heldt, 1985; Arrivault *et al*., 2017; Denton *et al*., 2017). We then measured ∼30 metabolites (including Tre6P) by LC-MS/MS, whereas pyruvate, PEP, 3PGA, DHAP, soluble sugars and starch were determined by enzymatic assays. With these data, we calculated the proportion of each metabolite in BSC and MC (Supplementary Tables S3-S5).

For Experiment 2, we estimated that ∼83% of Tre6P was in the BSC fraction (Fig. 5 and Supplementary Table S4). As already noted, some of this Tre6P might be in the phloem complex itself, rather than BSC. Other metabolites that were predominantly in the BSC fraction included pyruvate, 3PGA, sucrose and starch; conversely, DHAP was mainly found in the MC fraction, while PEP was almost equally distributed between BSC and MC (Fig. 5 and Supplementary Table S4). It should be noted that we estimated similar proportions for Tre6P, DHAP and starch in Experiment 1 (Supplementary Tables S3 and S5).

**Figure 5.**
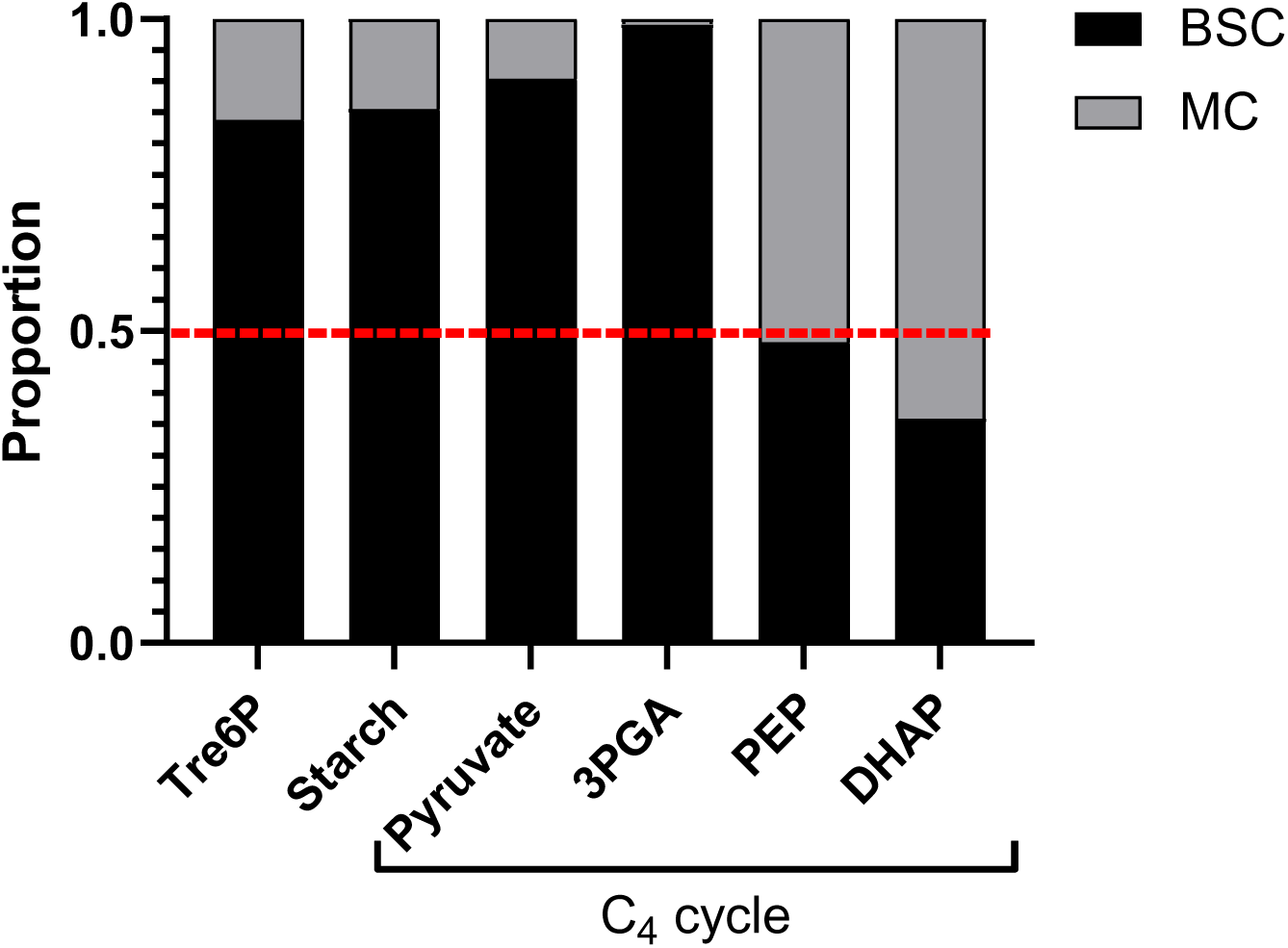
Distribution of selected metabolites in BSC and MC from *S. viridis* leaves. The abundances of each metabolite in BSC and MC were calculated as described in the methods section from metabolite/PRK *vs* PEPC/PRK plots (as shown in Supplementary Figure S6). Data are the mean of 4 biological replicates (except for starch, n=3) from one representative experiment (Experiment 2, Supplementary Table S4).

The levels of Tre6P and sucrose are tightly correlated across many treatments and conditions in *A. thaliana* leaves (Yadav *et al*., 2014). To test whether soluble sugars were correlated with Tre6P in *S. viridis* leaves, we measured these metabolites in whole leaves harvested every four hours during the light period, from Zeitgeber time (ZT) 0 to 16 h (Supplementary Fig. S11 and Supplementary Table S8). At the whole leaf level, Tre6P was not significantly correlated with glucose, fructose or sucrose (Fig. 6). As previously mentioned, the leaves of C_4_ plants have an additional complexity compared to those from C_3_ plants, due to the arrangement of MC and BSC in the characteristic Kranz anatomy. Thus, we performed the same analysis using data from the second fractionation experiment (Supplementary Fig. S11 and Supplementary Table S4). As shown in Fig. 6, across cell-type-enriched fractions, Tre6P was significantly correlated with sucrose and fructose and, to a lesser extent, with glucose.

**Figure 6.**
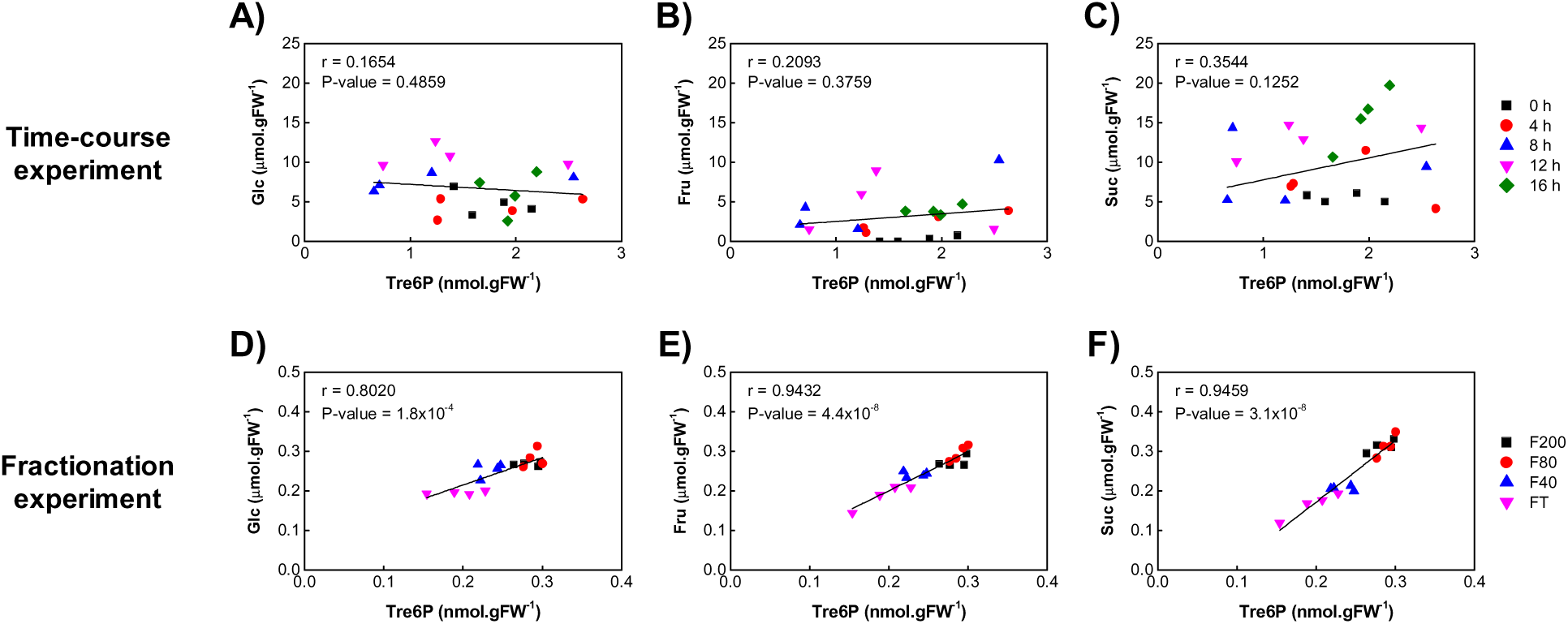
Correlation analysis of metabolites in fractions of *S. viridis* leaves. Data presented in panels A-C correspond to whole-leaf samples from the time-course experiment (Supplementary Table S8), while data presented in panels D-F correspond to one representative fractionation experiment (Experiment 2, Supplementary Table S4). Pearson correlation coefficients (r) between Tre6P and glucose (A and D), fructose (B and E) or sucrose (C and F) were calculated with a 95% confidence interval.

## Discussion

Tre6P metabolism is critical for plant growth and development (Fichtner and Lunn, 2021) and recent reports indicate that it might be an important metabolic target to engineer crops with improved agronomic traits (Kretzschmar *et al*., 2015; Nuccio *et al*., 2015; Griffiths *et al*., 2016; Gabriel *et al*., 2021). However, most of our knowledge on this signal molecule is based on studies performed with the model plant *A. thaliana*, a eudicot that performs C_3_ photosynthesis (Rojas *et al*., 2023). Here, we studied the compartmentation of Tre6P metabolism in *S. viridis*, a model system for studying C_4_ photosynthesis and associated traits (Brutnell *et al*., 2010; Doust *et al*., 2019).

To gain an overview of Tre6P and trehalose metabolism in *S. viridis*, we searched the genome sequence to identify all recognisable *TPS*, *TPP* and *TRE* genes, and then performed a phylogenetic analysis of the corresponding protein sequences. The number of genes found in this species (10 *TPS*, 10 *TPP* and 1 *TRE*) is the same as in its domesticated relative *S. italica* (Henry *et al*., 2020). The topologies of the reconstructed phylogenetic trees in this work (Fig. 1 and Supplementary Fig. S3) were in good agreement with those published by Henry *et al*. (2014) for Tre6P-related enzymes from maize, reinforcing the suitability of *S. viridis* species as a model system for photosynthesis research in crops.

The expression of transcripts encoding enzymes involved in Tre6P metabolism in *S. viridis* showed a differential pattern among the tissues analysed in this work. Analysis of Tre6P-related transcripts in different tissues of *S. viridis* was not previously reported, although other works showed their response in leaves subjected to different treatments, including light (Henry *et al*., 2014, 2020), salt (Henry *et al*., 2015), high light and temperature (Anderson *et al*., 2021). The *SvTPS1* transcript and SvTPS1 protein abundance was highest in internode samples (Supplementary Fig. S6 and Fig. 3D), which are rich in vascular tissue. This is in good agreement with data from *A. thaliana*, where the TPS1 protein was mainly detected in the vasculature of leaves, roots and reproductive tissues (Fichtner *et al*., 2020). We could not find Tre6P-related transcripts in the RNAseq analysis performed by Martin *et al*. (2016) using developing internodes from *S. viridis*, probably due to differences in the sampling procedures and analysis between both works. Other authors have reported that *A. thaliana TPS1* transcripts are relatively abundant in the protovasculature of developing leaves and the shoot apical meristem (Wahl *et al*., 2013), and in differentiated phloem cells from the inflorescence stem (Shi *et al*., 2021).

The transcripts encoding class II TPS proteins in *S. viridis* showed different levels according to the analysed tissue, which resembles data reported for class II TPS transcripts from *A. thaliana* (Ramon *et al*., 2009). As previously mentioned, our analyses were performed with samples harvested in the middle of the light period, and thus did not capture the diurnal fluctuations previously observed in *A. thaliana* (Usadel *et al*., 2008) and maize (Henry *et al*., 2014; Borba *et al*., 2023). The *SvTPS3* transcript displayed high levels in seedlings and whole leaves (Supplementary Fig. S6). The protein encoded by this transcript was located in a sub-clade exclusively containing class II TPS proteins from monocots, well separated from all other class II TPS sequences from eudicots (Fig. 1). This particular branch contains sequences from PACMAD (*S. viridis* and maize) and BOP/BEP (wheat, rice and *B. distachyon*) species, as well as from monocots outside the Poaceae that were included in our analysis (*A. comosus* and *M. acuminata*). This suggests that the monocot-specific class II TPS sub-clade arose before the divergence of the orders Poales (including the Poaceae and Bromeliaceae) and Zingeberales (including the Musaceae). The *SvTPS6* transcript was mainly detected in whole leaves and the fourth internode, while abundance of the *SvTPS7* transcript was highest in the flag leaf and the fourth internode (Supplementary Fig. S6). The proteins encoded by these transcripts were located in the same sub-clade as *A. thaliana* TPS7 (Fig. 1), whose expression was mainly detected in leaf primordia, cotyledons, roots (concentrated in the vasculature of the mature root and evenly distributed in the meristematic zone) and flowers (in ovules and pollen; Ramon *et al*., 2009). Recently, Van Leene *et al*. (2022) showed that all class II TPS proteins from *A. thaliana*, including TPS7, interact with and potentially act as regulators of the SUCROSE-NON-FERMENTING1-RELATED PROTEIN KINASE1 (SnRK1). Whether these proteins play the same role in *S. viridis* and other C_4_ grasses has to be further investigated.

Transcripts encoding TPP proteins from *S. viridis* also showed a particular distribution pattern, according to the tissue and cell type under analysis, as previously reported for *A. thaliana* (Vandesteene *et al*., 2012). The *SvTPP2* transcript was mainly detected in the fourth internode, with lower but significant levels in seedlings, whole leaves and the inflorescence (Supplementary Fig. S6). The protein encoded by this transcript is the orthologue of ZmTPP4; both were located in clade B, which also contains RAMOSA3 (ZmTPP10; Supplementary Fig. S3). The levels of *SvTPP1* were higher in whole leaves, the inflorescence, and the fourth internode, compared to other tissues, while levels of *SvTPP8* were higher in seedlings, the flag leaf and the fourth internode (Supplementary Fig. S6). The SvTPP1 and SvTPP8 protein sequences were located in clade A, which contains *A. thaliana* TPPA, TPPF and TPPG (Supplementary Fig. S3). There is evidence of recent duplication of genes encoding *A. thaliana* clade A TPPs, associated with whole-genome duplications within the Brassicaceae, with a high degree of paralogue retention and evidence of neofunctionalization (Vandesteene *et al*., 2012). Thus, it seems feasible that the duplication and diversification of clade A *S. viridis* TPPs observed in Supplementary Fig. S3 occurred independently of those in *A. thaliana* (Kerbler *et al*., 2023).

We found that several Tre6P-related transcripts were preferentially located in BSC of *S. viridis* leaves (Fig. 2). To compare our results with previously published datasets, we retrieved RNAseq data from studies performed with enriched MC and BSC fractions from *S. viridis* and maize (John *et al*., 2014; Denton *et al*., 2017; Supplementary Fig. S12 and Supplementary Table S9). Our analysis showed that the *TPS1* transcript was enriched in BSC from *S. viridis* and maize leaves (Fig. 2B and Supplementary Fig. S12), as previously shown for the *A. thaliana TPS1* transcript (Aubry *et al*., 2014). A similar analysis was performed by Benning *et al*. (2023), who used publicly available RNA deep-sequencing data to map the distribution of sugar sensing genes in leaves of C_3_ and C_4_ species; among these, they found that *TPS1* transcripts from C_4_ species were mainly located in BSC. It is important to recall that our experimental data originate from samples harvested in the middle of the day (ZT 8 h), as *TPS* and *TPP* transcripts fluctuate during the diel cycle (Usadel *et al*., 2008, Borba *et al*., 2023); particularly, the maize *TPS1* transcript displayed the highest level at ZT 8 h (Borba *et al*., 2023), indicating that this time of day is optimal for analysing the cellular distribution of *TPS1* transcripts. Regarding the class II *TPS* transcripts, the levels of *SvTPS3* and *SvTPS10* were significantly higher in BSC than in MC when analysed by RT-qPCR (Fig. 2B); however, these patterns were not the same in RNAseq data from *S. viridis* and maize. As shown in Supplementary Fig. S12, *SvTPS9* and *SvTPS10* displayed the highest enrichment in *S. viridis* BSC, whereas *ZmTPS11* (which clustered together with *SvTPS7*; Fig. 1) showed high enrichment in BSC from almost all leaf segments of maize leaves. Abundance of *SvTPP1* (the orthologue of *ZmTPP6*) was similar in MC and BSC (Fig. 2C), which agrees with RNAseq data from *S. viridis* and maize (Supplementary Fig. S12). Our RT-qPCR results showed greater abundance of *SvTPP2, SvTPP3*, *SvTPP7* and *SvTPP9* in BSC than in MC (Fig. 2C), and the same trend was observed in the RNAseq data from *S. viridis* (Supplementary Fig. S12). Similarly, *ZmTPP1*, *ZmTPP2*, *ZmTPP7* and *ZmTPP8* (the orthologues of *SvTPP7*, *SvTPP3*, *SvTPP2* and *SvTPP3*, respectively; Supplementary Fig. S3) were enriched in BSC across all fractions of the maize leaf, while *ZmTPP4* and *ZmTPP9* (the orthologues of *SvTPP2* and *SvTPP5*, respectively; Supplementary Fig. S3) were enriched in BSC from the middle and the tip of the maize leaf (fractions 1 to 3, Supplementary Fig. S12). The levels of the *SvTRE* transcript were similar in MC and BSC, as reported in RNAseq studies of *S. viridis* and maize leaves (Fig. 2D and Supplementary Fig. S12).

Cell-specific expression of transcripts in MC and BSC is regulated by transcriptional and post-transcriptional mechanisms (Reeves *et al*., 2017). Furthermore, due to translational and post-translational regulation, transcript abundance is not necessarily a reliable indicator of protein abundance (Gibon *et al*., 2004; Wang *et al*., 2011; Reeves *et al*., 2017). Thus, we investigated the distribution of the SvTPS1 polypeptide in different tissues and cell types. Immunoblotting assays detected the SvTPS1 protein in whole leaves and the fourth internode (Fig. 3D), as previously observed in the transcript abundance analysis (Supplementary Fig. S6). The localization of SvTPS1 in the BSC fraction of *S. viridis* leaves by immunoblotting (Fig. 3B-C) is consistent with the proteomic data from Calace *et al*. (2021), presented in Fig. 3A.

The antiserum raised against SvTPS1 consistently recognized two immunoreactive protein bands, particularly in BSC (Fig. 3B-C). The identity of the protein detected by the anti-SvTPS1 was confirmed by immunoprecipitation and mass spectrometry (Supplementary Fig. S8). Based on our results, we hypothesized that these proteoforms of SvTPS1 may result from two splicing variants or a post-translational modification. The *S. viridis* genome sequence has only one annotated gene model for *TPS1* (*Sevir.3G180900.1*), whereas *S. italica* has two gene models for *TPS1* (*Seita.3G176900.1* and *Seita.3G176900.2*), in which the alternative combination of the first two exons could produce proteins with different N-terminus, with predicted molecular masses of 108.4 and 94.9 kDa, respectively (Supplementary Fig. S13). The *Seita.3G176900.1* and *Seita.3G176900.2* transcripts displayed a differential response to light (Henry *et al*., 2020), suggesting their encoded polypeptides could have particular biological functions. A detailed comparison of the *Sevir.3G180900* and *Seita.3G176900* DNA sequences showed eight differences, but only from the fifth intron towards the 3’-end (Supplementary Fig. S13). Therefore, an alternative transcript could also be produced from the *Sevir.3G180900* gene. Regarding the existence of a putative post-translational modification, it is important to note that *A. thaliana* TPS1 is phosphorylated and potentially sumoylated (Fichtner *et al*., 2020; Fichtner and Lunn, 2021). The phosphorylation sites found on the *A. thaliana* TPS1 are not conserved in SvTPS1 but the putative sumoylation site (Lys^902^ in *A. thaliana* TPS1, Lys^919^ in SvTPS1) is well-conserved (Supplementary Fig. S4; Fichtner *et al*., 2020; Fichtner and Lunn, 2021). Considering the theoretical molecular mass of SvTPS1 (108.4 kDa) and the estimated sizes of the immunoreactive bands (112 and 118 kDa, Fig. 3B-C), the occurrence of a post-translational modification seems more plausible, as the two putative splicing variants of SvTPS1 should have 108.4 and 94.9 kDa, respectively (Supplementary Fig. S13).

To confirm the immunoblotting results obtained with enriched fractions, we performed immunohistochemistry on cross-sections of *S. viridis* leaves. SvTPS1 was preferentially detected in BSC (Fig. 4 and Supplementary Fig. S9). We did not detect any significant signal in the vascular bundle, suggesting SvTPS1 is not present in this compartment. This finding represents a major difference with the observation made for *A. thaliana* TPS1, which was primarily detected in the companion cell-sieve element complex and, to a lesser extent, in the phloem parenchyma (BSC) and xylem parenchyma cells (Fichtner *et al*., 2020). Overall, our results show that the only class I TPS from *S. viridis* is predominantly located in BSC, suggesting this is the major location for Tre6P synthesis in *S. viridis* leaves.

We adapted a method, originally developed for maize, that allows metabolite analysis in MC and BSC enriched fractions (Stitt and Heldt, 1985). We found that the proportion of Tre6P was higher in BSC than in MC. Similar results were observed for pyruvate, 3PGA and starch, as expected for an NADP-ME C_4_ species (Fig. 5 and Supplementary Tables S3-S5). Similarly, ribulose 1,5-bisphosphate and other intermediates of the Calvin-Benson cycle were almost exclusively located in BSC (Supplementary Table S5). The amount of malate was also higher in BSC than MC. This result is counterintuitive, as the concentration gradient of malate is expected to be the opposite; i.e., a higher concentration in MC than BSC, to drive diffusion of malate from the MC to the BSC. However, it is important to consider that maize leaves have a large pool of malate (up to 60% of the total malate) that is not actively involved in the C_4_ cycle, including a potentially large pool in the vacuole (Arrivault *et al*., 2017). Thus, there still might be a concentration gradient which could move malate from the cytosol of MC to cytosol of BSC in *S. viridis* leaves, although this has to be proven, for instance, by a combination of ^13^CO_2_ labelling and leaf fractionation, as previously done with maize leaves (Arrivault *et al*., 2017). Two intermediates of the Calvin-Benson cycle showed contrasting distributions: 3PGA was predominantly located in the BSC, while DHAP was mainly in the MC (Fig. 5). This is similar to the intercellular distributions of 3PGA and DHAP reported for maize leaves (Leegood, 1985; Stitt and Heldt, 1985; Lunn and Furbank, 1997; Arrivault *et al*., 2017), where there is intercellular shuttling of these metabolites. Pyruvate was also predominantly located in the BSC, consistent with its production from malate decarboxylation, via NADP-ME, in the BSCs, and diffusion into the MCs for conversion to PEP via PPDK. PEP was evenly distributed between MC and BSC (Fig. 5 and Supplementary Table S5). These distributions differ from those reported for maize, where PEP showed a preferential increase in the BSC and pyruvate was located mainly in the MC (Arrivault *et al*., 2017). Such differences could be explained by i) the existence of metabolic gradients along the leaf, or ii) the presence of metabolite pools that are not actively involved in C_4_ photosynthesis, for example, if located in the vacuole, which could mask metabolite gradients between the cytosol of MC and that of BSC (Szecowka *et al*., 2013; Arrivault *et al*., 2017; Denton *et al*., 2017).

A limitation of our study is that samples for metabolite analysis were collected from the middle section of the leaf blade at one defined time-point in the middle of the light period, so we could not capture metabolic dynamics. It would be interesting to perform labelling studies followed by cellular enrichment to obtain a detailed scenario of the metabolic fluctuations that occur in *S. viridis* leaves. Furthermore, overall gradients between the MC and BSC might not reflect the gradients between the cytosol of these two cell types, if there were active transport of metabolites across the plastid envelope or the pH gradient across the envelope membrane influences their distribution between the chloroplast stroma and the cytosol, as is known to be the case for 3PGA (Fliege *et al*., 1978; Stitt *et al*., 1980). The relation between the overall intercellular metabolite gradient and the gradient between the cytosols of the BSC and MC will also be affected by the relative contribution of chloroplasts and cytosol to cytoplasmic volume. Overall, our study indicates that the metabolite gradients reported in maize may not be generalizable and underlines the need for studies of metabolite gradients in other C_4_ species. Just as different biochemical solutions have arisen in different species to concentrate CO_2_ into the BSC (Furbank and Kelly, 2021), different strategies may also have been followed to generate the intercellular gradients in the cytosol that are needed to drive metabolite movement between the BSC and MC. Overall, our fractionation data provide the first insights into the distributions of key photosynthetic intermediates in *S. viridis*. These are consistent with the photosynthetic fluxes and intercellular metabolite movements that we would expect to see in an NADP-ME-type C_4_ species, demonstrating the reliability of this experimental approach.

The main aim of the cell fractionation experiments in our study was to determine the cellular localization of Tre6P. Our transcript and protein analyses showed that the SvTPS1 enzyme is predominantly located in the BSC and associated vascular tissues, and essentially absent from MC (Figs 2-4), indicating that Tre6P is synthesized in BSC but not in MC. We found that approximately 83% of Tre6P was present in the BSC fraction (Fig. 5) and, considering the previously mentioned limitations, this value could be even higher. It is worth noting that some of the Tre6P in the BSC fraction may be in the phloem complex because, although the cell separation method allows the sample to be enriched with BSC strands, this enrichment is partial and may bring with it remains from other cells types. This would make sense considering that, in *A. thaliana*, Tre6P metabolism has been reported to take place in both phloem parenchyma (equivalent to the BSC) and in the companion cell-sieve element complex of the phloem (Fichtner *et al*., 2020). As previously mentioned, sucrose synthesis seems to occur in MC of *S. viridis* leaves; however, the amount of sucrose was higher in the BSC fraction than in the MC fraction (Supplementary Table S5). This result is puzzling, as sucrose levels are expected to be higher in the MC than the BSC, to drive its diffusion to the BSC. We thus speculate that i) sucrose is sequestered either in some compartment other than the cytosol in the BSC or other cell types that are enriched in the BSC fractions; or ii) that a large part of the sucrose in the leaf is in the phloem.

Lunn *et al*. (2006) showed there is a significant positive correlation between Tre6P and sucrose levels in samples from *A. thaliana* seedlings and rosettes. The correlation between Tre6P and sucrose persisted even in transgenic lines expressing the *E. coli* TPS (otsA) and TPP (otsB) proteins under the control of the *35S* promoter, although sucrose levels were markedly lower or higher, respectively, which led to the development of the Tre6P-sucrose nexus model (Yadav *et al*., 2014). Tre6P and sucrose levels fluctuated during the light period in *S. viridis* leaves (Supplementary Fig. S11) but, unlike *A. thaliana*, we did not observe a significant correlation at the whole-leaf level (Fig. 6). Conversely, we found a significant correlation between Tre6P and sucrose across leaf fractions containing different proportions of MC and BSC (Fig. 6). The Tre6P-sucrose nexus model postulates that Tre6P is a signal and a negative regulator of sucrose levels (Yadav *et al*., 2014). According to this model, if sucrose levels rise too high, Tre6P rises in parallel and triggers mechanisms to lower the level of sucrose, either by reducing sucrose synthesis or increasing its degradation or transport. Regarding the first alternative, research performed with ethanol-inducible *A. thaliana* transgenic lines showed that artificially increased Tre6P levels decreased the amount of sucrose by diverting newly fixed carbon to organic and amino acids during the day, through the post-translational activation of PEPC and nitrate reductase (Figueroa *et al*., 2016). As previously mentioned, the initial CO_2_ fixation step in C_4_ plants is catalysed by PEPC, which is located in MC. Our results strongly suggest that Tre6P is mainly produced and accumulated in BSC of *S. viridis* leaves. Thus, if the Tre6P-PEPC connection is operational in C_4_ species, this would imply that Tre6P has to diffuse from BSC to MC; indeed, a small amount of Tre6P was actually found in MC (Fig. 5). The asymmetrical distribution of Tre6P suggests that it could be sensing sucrose levels in BSC and then act as a signal that moves into the MC to potentially modulate PEPC activity and/or sucrose synthesis, which is also located in MC. The molecular mechanism underlying the putative transport of Tre6P from BSC to MC deserves to be further studied but is beyond the scope of this work.

Another possibility is that Tre6P regulate sucrose levels by controlling phloem loading. TPS1 has been found in the vasculature of *A. thaliana* leaves; particularly, in the phloem parenchyma and the companion cell/sieve element complex (Fichtner *et al*., 2020). Tre6P has been shown to modulate the levels of *ZmSWEET13a,c* and *AtSWEET11-14* transcripts in maize and *A. thaliana*, respectively, which encode sucrose transporters involved in phloem loading (Oszvald *et al*., 2018; Fichtner *et al*., 2021). Single-cell transcriptomics in *A. thaliana*, rice and maize showed that *SWEET* transcripts were preferentially detected in BSC (Bezrutczyk *et al*., 2021; Shi *et al.,* 2021; Hua *et al.,* 2021; Tao *et al*., 2022). Moreover, the *ZmSWEET13a-c* transcripts and the SWEET13a protein were specifically detected in the abaxial BSC of major and intermediate veins from maize leaves (Bezrutczyk *et al*., 2021), while both SWEET13a and SWEET13b proteins were detected in the abaxial BSC of intermediate veins from *S. viridis* leaves (Chen *et al*., 2022). Based on these data, it has been proposed that sucrose moves from adaxial to abaxial cells via plasmodesmata and is then exported by SWEET13 to the apoplast, from where it is taken up by transporters from the SUT/SUC (sucrose-H^+^ symporter) family located in the companion cells (Bezrutczyk *et al*., 2021; Chen *et al*., 2022). The maize *zmsweet13a-c* triple mutant is impaired in sugar phloem loading, thus producing drastic effects on growth (Bezrutczyk *et al*., 2018). In *S. viridis*, such mutants have not yet been identified, but there is compelling evidence that it could be an apoplastic phloem-loading species, and thus SvSWEET13a-b might function in phloem loading (Chen *et al*., 2022). Based on this evidence, and the fact that Tre6P metabolism is mostly located in BSC, we speculate that Tre6P modulates sucrose export in leaves of C_4_ plants by controlling the expression, and thereby maximal activity, of the SWEET13 transporters, although this hypothesis remains to be tested. If confirmed, this would implicate Tre6P in the reconfiguring of phloem loading that is a common feature of C_4_ monocots, and potentially an essential element in the evolution of Kranz anatomy and C_4_ photosynthesis (Furbank and Kelly, 2021; Rojas *et al*., 2023).

To conclude, our results indicate that Tre6P is synthesized and mainly located in the BSC of *S. viridis*, which play a central role in phloem loading in C_4_ species and represent a strategic site at the interface between source and sink tissues. This localization would allow Tre6P to monitor the levels of sucrose in BSC and, via diffusion into the MC, potentially regulate CO_2_ fixation and sucrose biosynthesis as well as sucrose export from the leaves (Rojas *et al*., 2023).

## Supplementary Data

**Supplementary File S1.** Optimised DNA sequence for the recombinant expression of SvTPS1 in *E. coli* cells.

**Supplementary Figure S1.** Analysis of MC and BSC fractions for transcript and protein determinations.

**Supplementary Figure S2.** C_t_ values for the reference transcript *SvKIN* in different tissues and cell types from *S. viridis*.

**Supplementary Figure S3.** Phylogenetic relationships of *Setaria viridis* TPP proteins.

**Supplementary Figure S4.** Multiple sequence alignment of *S. viridis* TPS proteins.

**Supplementary Figure S5.** Multiple sequence alignment of *S. viridis* TPP proteins.

**Supplementary Figure S6.** Relative abundance of transcripts encoding Tre6P-related enzymes in different *S. viridis* tissues.

**Supplementary Figure S7.** Immunoblotting of SvTPS1 in different tissues and cell types of *S. viridis*.

**Supplementary Figure S8.** Analysis by mass spectrometry of immunoprecipitated SvTPS1.

**Supplementary Figure S9.** Immunolocalization of SvTPS1 in cross-sections of *S. viridis* leaves.

**Supplementary Figure S10.** Activities of marker enzymes in different fractions from *S. viridis* leaves.

**Supplementary Figure S11.** Analysis of metabolites from *S. viridis* leaves.

**Supplementary Figure S12.** Analysis of transcripts encoding Tre6P-related enzymes in BSC and MC from *S. viridis* and maize leaves.

**Supplementary Figure S13.** Gene models for the class I TPS from *S. viridis* and *S. italica*.

**Supplementary Table S1.** TPS and TPP protein sequences used for reconstruction of phylogenetic trees.

**Supplementary Table S2.** Sequences of the primers used to measure transcript levels by qPCR.

**Supplementary Table S3.** Metabolite and enzyme data and calculations (Experiment 1).

**Supplementary Table S4.** Metabolite and enzyme data and calculations (Experiment 2).

**Supplementary Table S5.** Calculation of metabolite fractions in MC and BSC.

**Supplementary Table S6.** Analysis of immunoprecipitated SvTPS1 by mass spectrometry.

**Supplementary Table S7.** Recovery of marker enzymes in the fractionation experiments.

**Supplementary Table S8.** Metabolite data from *S. viridis* leaves during a day time-course.

**Supplementary Table S9.** Relative transcript abundance in MC and BSC from *S. viridis* and maize.

## Acknowledgements

TT, BER and PC are fellows from CONICET. CS is a fellow from Agencia I+D+i. LEL, MSaigo and CMF are researchers from CONICET. CMF thanks NME, CF and MF for their everyday support. We thank to the Mass Spectrometry Unit from IBR (UNR-CONICET) for the analysis of immunoprecipitated SvTPS1; Agustín Arce for help with statistical analysis; Andrés Dekanty for providing the Cy2-conjugated goat anti-rabbit IgG; and Sergio Gonzalez for technical assistance with the confocal microscope.

## Author contributions

Conceptualization: MStitt, JEL and CMF; Investigation: TT, BER, RF, CS, LEL, JVC, PC, SA and CMF; Formal analysis: all authors; Writing - Original Draft: TT, BER and CMF; Writing - Review & Editing: all authors.

## Data availability

All data supporting the findings of this study are available within the paper and within its supplementary data published online. The raw MS data resulting from the analysis of the immunoprecipitated TPS1 were deposited to the PRIDE repository with the dataset identifier PXD057139.

## Abbreviations

BSC: bundle sheath cells
MC: mesophyll cells
NADP-MDH: NADP-dependent malate dehydrogenase
NADP-ME: NADP-dependent malic enzyme
PEPC: phospho*enol*pyruvate carboxylase
PRK: phosphoribulokinase
SWEET: SUGARS WILL EVENTUALLY BE EXPORTED TRANSPORTERS
Tre6P: trehalose 6-phosphate
TPS: Tre6P synthase
TPP: Tre6P phosphatase
TRE: trehalase.

