## Supplementary Figures for "Intercellular compartmentation of trehalose 6-phosphate metabolism in *Setaria viridis* leaves"

^#^Equal contribution

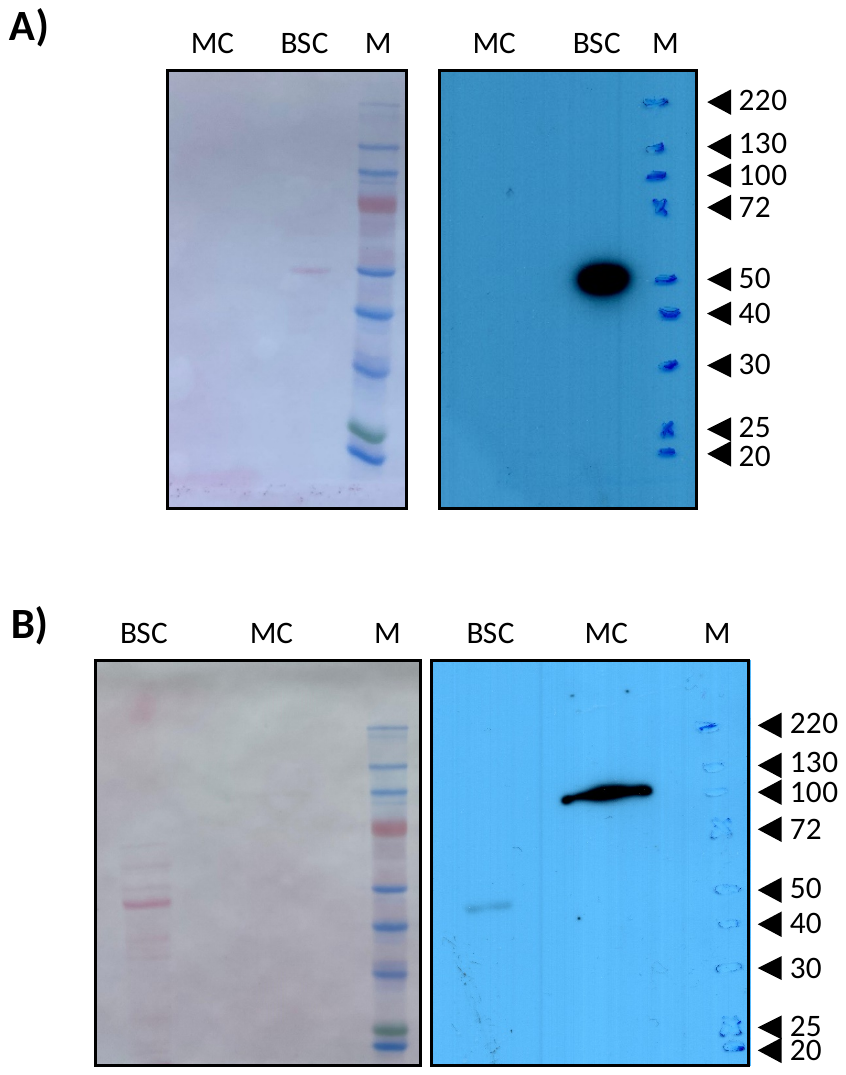

**Supplementary Figure S1. Analysis of MC and BSC fractions for transcript and protein determinations.** (A) Immunoblotting of the Rubisco large subunit (RbcL). The amount of protein loaded were 10 µg for MC and 1 µg for BSC. The anti-RbcL antibody was diluted 1:10,000 and the membrane was incubated at room temperature for 90 min. (B) Immunoblotting of PEPC. The amount of protein loaded were 10 µg for both MC and BSC. The anti-PEPC antibody was diluted 1:1,000 and the membrane was incubated at room temperature for 90 min. BSC, protein extract from the fraction enriched in bundle sheath cells; MC, protein extract from the fraction enriched in mesophyll cells; M, prestained molecular mass marker. Numbers on the right indicate the mass of the markers in kDa. Left panels, loading controls (membranes stained with Ponceau Red); right panels, western blots (X-ray films revealed with ECL).

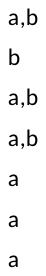

**Supplementary Figure S2. C_t_ values for the reference transcript *SvKIN* in different tissues and cell types from *S. viridis*.** IF; inflorescence, FL; flag leaf, IN; internode, SL; seedlings, MC; mesophyll cells, BSC; bundle sheath cells, WL; whole leaf. Data are the mean ± standard error of three biological replicates. Different small-case letters indicate statistically significant differences according to the Tukey test with a 95% confidence.

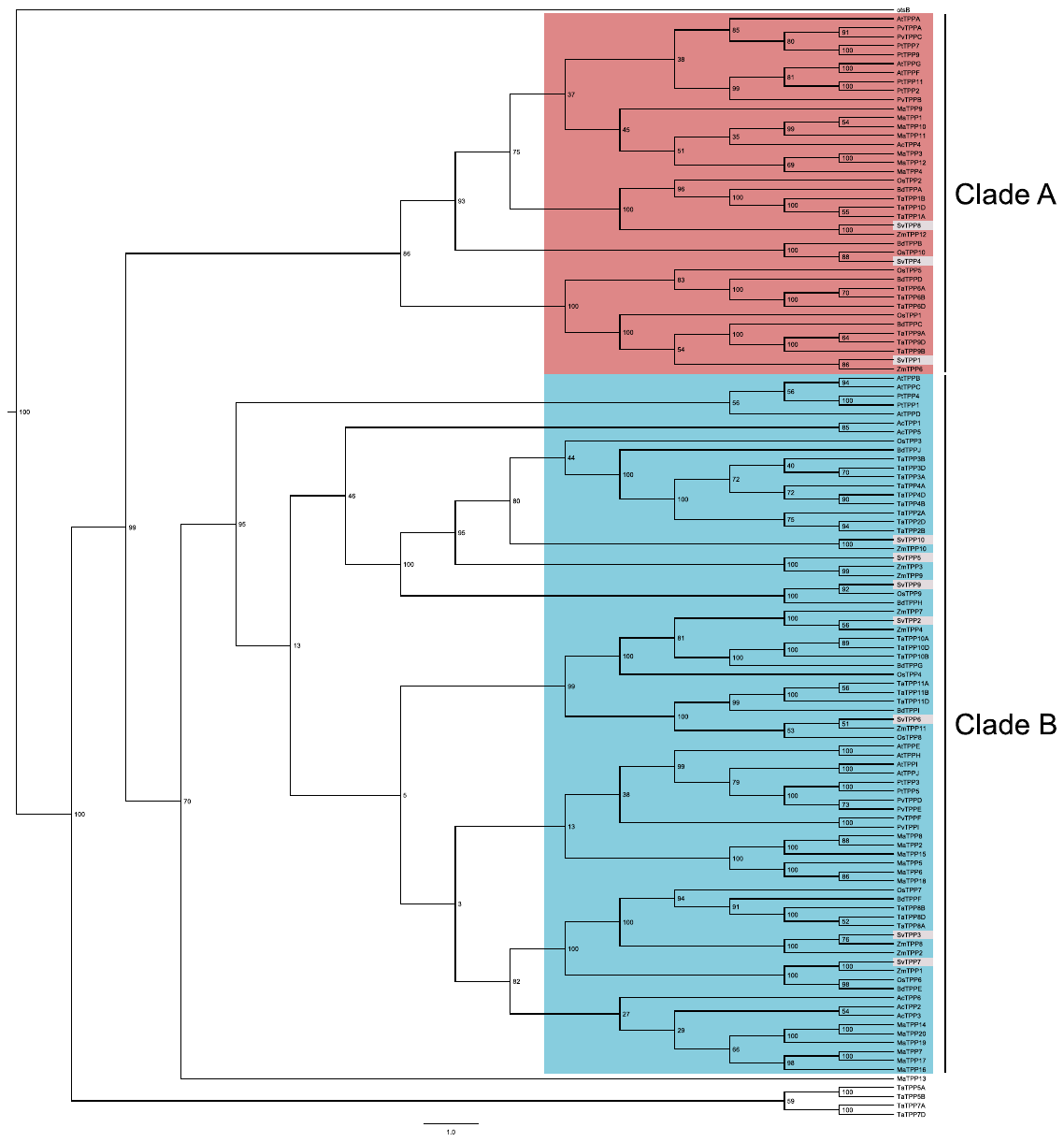

**Supplementary Figure S3. Phylogenetic relationships of *Setaria viridis* TPP proteins.** TPP protein sequences were collated from *Setaria viridis* (Sv) and other selected species: Ac, *Ananas comosus*; At, *Arabidopsis thaliana*; Bd, *Brachypodium* *dystachion*; Mc, *Musa acuminata*; Os, *Oryza sativa*; Pt, *Populus trichocarpa*; Pv, *Phaseolus vulgaris*; Ta, *Triticum aestivum*; Zm, *Zea mays* (see Supplementary Table S1 for gene/protein identifiers). A distance-based tree was reconstructed using the BioNJ algorithm. Number in nodes represent the bootstrap obtained for 1000 replicates. The tree is rooted on the *Escherichia coli* TPP (otsB). Sub-clades in the tree are coloured according to the position of *A. thaliana* TPP sequences: AtTPPA/F/G (red) and AtTPPB/C/D/E/H/I/J (blue). *S. viridis* sequences are highlighted in grey.

```
                        10         20         30         40         50         60         70         80         90        100
               ....|....| ....|....| ....|....| ....|....| ....|....| ....|....| ....|....| ....|....| ....|....| ....|....|
otsA           ---------- ---------- ---------- ---------- ---------- ---------- ---------- ---------- ---------- ----------
AtTPS1         ---------- MPGNKYNCSS SHIPLS---- ---------- ---------- ---RTERLLR DRELREKRKS NRARNPNDVA GSSENSENDL RLEGD--SSR
SvTPS1         MSSDAAGGQR SSRNRGRADA APMPTSSPFT GDGGGAGSP- ---------- --TRVERMLR EREHSRRHIF ASDTMDTDAA EPVFASAGAF AADGVQ-SPG
SvTPS2         ---------- ---------- ---------- ---------- ---------- ---------- ---------- ---------- ---------- ----------
SvTPS3         ---------- MPSLSC---- --HNLLDLAA ADEVPL---- ---------- PSPTPLRLPR ---------- -VMSVASPA- ---------- -----SP---
SvTPS4         ---------- MVLNSF---- --SNLLDICS EDVFDF---- ---------- -QQPLRSLPC ---------- -AVTSPGIRS DPDWESSNGS NLIGS-----
SvTPS5         ---------- MVSRSY---- --SNLLELAS GG-------- -GGGGGGEPL PSLGRRRIPR ---------- -VVTASGIVP DLDASDAD-- ADAASAA---
SvTPS6         ---------- MFSRSY---- --TNLLDLAN GNLSALDYGG GGGGGGGGGG RPPRARRMQR ---------- -TMTTPGTLV ELDEER---- --AGSVA---
SvTPS7         ---------- MMSRSY---- --TNLLDLAE GNFAALGPAA GGGRRRQG-- -SFGTKRMSR ---------- -VMTVPGTLS ELDGEDDSEP AATNSVA---
SvTPS8         ---------- MVSKSY---- --SNLLDMTS GDGFDF---- ---------- -RQPFKSLPR ---------- -VVTSPGIIS DPDWDTRSDG DSVGS-----
SvTPS9         ---------- MPSLSC---- --HNLLDLAA ADEVPL---- ---------- PSPTPLRLPR ---------- -VMSVASPA- ---------- -----SP---
SvTPS10        ---------- MVSRSY---- --SNLLDLAT GA-------- -ADQAPAPTA LGALRRRLPR ---------- -VVTNTGLID DSPAS----- ----------

                       110        120        130        140        150        160        170        180        190        200
               ....|....| ....|....| ....|....| ....|....| ....|....| ....|....| ....|....| ....|....| ....|....| ....|....|
otsA           ---------- ---------- ---------- ---------M SRLVVVSNRI APPD------ --EHAASAGG LAVGILGAL- ---------K AAGGLWFGWS
AtTPS1         QYVEQYLEGA AAAMAHDDAC ERQEVRPYN- ---------R QRLLVVANRL PVSAVRRG-- --EDSWSLEI SAGGLVSALL -------GVK EFEARWIGWA
SvTPS1         RASPANMEDA GG-AASGHAA RPPLAGSRSG FRRLGLRGMK QRLLVVANRL PVSANRRG-- --EDQWSLEI SAGGLVSALL -------GVK DVDAKWIGWA
SvTPS2         ---------- ---------- ---------- ---------- ---------- ---------- ---------- ---------- ---------- ----------
SvTPS3         ---------- ---------- ---------- -TSP---PTP PRRVIVSHRL PLRAAPDP-- AAPFGFAFTV DAGTVAYQLR SGL-----PA SAPVLHIGTL
SvTPS4         ---------- ---------- ---------- -----APPCL TRKIVVANFL PLNCTKDE-- -ATREWSFAV DDNQLLVQLK DGF-----PI DSEVIYVGSL
SvTPS5         ---------- ---------- ---------- -SDHSSHAPR ERVIIVANQL PVRATRRAGP GTGAGWDFAW DEDSLLLQVK DSLRAHHGRA DVEFVYVGGL
SvTPS6         ---------- ---------- ---------- -SDVQSSLAG DRLIVVANTL PVRGERRP-- -DGRGWTFSW DEDSLLFHLR DGL-----PD DMEVLYVGSL
SvTPS7         ---------- ---------- ---------- -SDVPSSLAA DRMIVVSNQL PVVARRRP-- -DGRGWSFSW DDDSLLLQLR DGI-----PD EMEVLFVGSL
SvTPS8         ---------- ---------- ---------- -----AS-FS ERKIIVANFL PLNCTRDE-- -AG-QLSFSL DDDALLVQLK HGF-----SN ETDVVYVGSL
SvTPS9         ---------- ---------- ---------- -TSSPSPPAT PRRVIVSHRL PLHAAPDP-- AAPFGFAFTV DAGTVAYQLR SGL-----PA SAPVLHIGTL
SvTPS10        ---------- ---------- ---------- -PSTPSPAPR PRTIVVANQL PIRSHRPASP E--EPWTFDW DEDSLLRHLH -----HTSPP SMEFIYIGCL

                       210        220        230        240        250        260        270        280        290        300
               ....|....| ....|....| ....|....| ....|....| ....|....| ....|....| ....|....| ....|....| ....|....| ....|....|
otsA           GETG-NEDQP LKKVKKGNIT ---WASFNLS EQDLDEYYNQ FSNAVLWPAF HYRLD----- ---------- ---LVQFQ-- RPAWDGYLRV NALLADKLLP
AtTPS1         GVNVP-DEVG QKALSKALAE KR-CIPVFLD EEIVHQYYNG YCNNILWPLF HYLGLP---- -------QED --RLATTRSF QSQFAAYKKA NQMFADVVNE
SvTPS1         GVNVP-DEVG QRALTRALAE KR-CIPVFLD EEIVHQYYNG YCNNILWPLF HYLGLP---- -------QED --RLATTRNF ESQFDAYKRA NQMFADVVYQ
SvTPS2         ---------- --------ME NFSCLPVHLD GGRHAEFYDG FCKRYLWPLL HYLLPFTVA- ---------- DAGNLYFD-- EAEYRAFIAA NRQFADRVIQ
SvTPS3         PAAAA--EAA SDELSNYLLA NFSCLPVYLP SDLHHRFYHG FCKHYLWPLL HYLLPLTPS- ---------- SLGGLPFQ-- RTLYHSFLSA NRAFADRLTE
SvTPS4         NVQVD--PSE QDQVSQKLFK EHKCIPTFLP ADLQQQFYHS FCKQHLWPLF HYMLPVCHD- ---------- --KGELFD-- RSLFQAYVRA NQIFADKVME
SvTPS5         RDDVP--PAD HDEVAHDLLE GFRCVPTFLP ADLRSRFYHG FCKQQLWPLF HYMLPLSPE- ---------- --LGGRFD-- RQLWQAYVSV NKIFADKILE
SvTPS6         RADVP--PAE QDDVAQALLE RFRCVPAFLP KDICDRFYHG FCKQMLWPLF HYMLPFSPD- ---------- --HGGRFD-- RSQWEAYVLT NKLFSQRVIE
SvTPS7         RADVP--VAE QDEVSQALLD RFRCAPVFLP DHLNDRFYHG FCKRQLWPLF HYMLPFSSSA SAATTSS--- SPGNGRFD-- RSAWEAYVLA NKFFFEKVVE
SvTPS8         KIQVD--PSE QDQVAQKLLR EYRCIPTFLP SDLQQQFYHG FCKQQLWPLF HYMLPICLD- ---------- --KGELFD-- RSLFQAYVRA NKLFADKVME
SvTPS9         PAAAA--EAA SDELSNYLLA NFSCLPVYLP SDLHHRFYHG FCKHYLWPLL HYLLPLTPS- ---------- SLGGLPFQ-- RTLYHSFLSA NRAFADRLTE
SvTPS10        RDDIP--AAD QDAVAQALLD THNCVPAFLP PDIAERYYHG FCKQHLWPLF HYMLPLSPD- ---------- --LGGRFD-- RSLWQAYVSA NKIFADKVLE

                       310        320        330        340        350        360        370        380        390        400
               ....|....| ....|....| ....|....| ....|....| ....|....| ....|....| ....|....| ....|....| ....|....| ....|....|
otsA           LLQDDD--II WIHDYHLLPF AHELRKRGVN -NRIGFFLHI PFPTPEIFNA LPTYDTLLEQ LCDYDLLGFQ TENDRLAFLD CLSNLTRVTT RS------AK
AtTPS1         HY--EEGDVV WCHDYHLMFL PKCLKEYNS- KMKVGWFLHT PFPSSEIHRT LPSRSELLRS VLAADLVGFH TYDYARHFVS ACTRILGLEG TP-------E
SvTPS1         HY--QEGDVI WCHDYHLMFL PKCLKDHDN- NMKVGWFLHT PFPSSEIYRT LPSRLELLRS VLCADLVGFH TYDYARHFVS ACTRILGLEG TP-------E
SvTPS2         VLSPDDGDLV IVHDYHLWAL PTFLRRKCPR -AAVGLFLHT PFPSAEILSS VAVCDELLRG LLNADLVGFH TSYYAHQFVS CCWGLLGVSY NISSLHGGHV
SvTPS3         VLSPDE-DLV WIHDYHLLAL PTFLRKRFPR -AKVGFFLHS PFPSSEIFRT IPVRDDLVRA LLNADLVGFH TFDYARHFLS ACSRLLGLDY QS---KRGYI
SvTPS4         AVNSDD-DCV WVHDYHLMLV PTFLRKKLHR -IKVGFFLHS PFPSSEIYKT LPVRDEILKS LLNADLIGFQ TFDYARHFLS CCSRLLGLNY ES---KRGHI
SvTPS5         VISPDE-DFV WVHDYHLMVL PTFLRKRFNR -VKLGFFLHS PFPSSEIYKT LPVREELLRS LLNADLIGFH TFDYARHFLS CCSRMLGLKY ES---QRGYI
SvTPS6         VLNPED-DYV WIHDYHLLAL PSFLRRRFNR -LRIGFFLHS PFPSSELYRS LPVRDEILKS LLNCDLIGFH TFDYARHFLS CCSRMLGIEY QS---KRGYI
SvTPS7         VINPED-DYV WVHDYHLMAL PTFLRRRFNR -LRIGFFLHS PFPSSEIYRT LPVREEILKA LLNCDLIGFH TFDYARHFLS CCSRMLGIEY QS---KRGYI
SvTPS8         VINTDD-DYV WVHDYHLMLL PTFLRKRLHR -IKLGFFLHS PFPSSEIYRT LPVRDEILKS LLNADLIGFQ TFDYARHFLS CCSRLLGLHY ES---KRGYI
SvTPS9         VLNPDE-DLV WIHDYHLLAL PTFLRKRFPR -AKVGFFLHS PFPSSEIFRT IPVRDDLVRA LLNADLVGFH TFDYARHFLS ACSRLLGLDY QS---KRGYI
SvTPS10        VINPDD-DFV WVHDYHLMVL PTFLRKRFNR -IKLGFFLHS PFPSSEIYKT LPVREELLRA LLNSDLIGFH TFDYARHFLS CCGRMLGLSY ES---KRGHI

                       410        420        430        440        450        460        470        480        490        500
               ....|....| ....|....| ....|....| ....|....| ....|....| ....|....| ....|....| ....|....| ....|....| ....|....|
otsA           SHTAWGKAFR TEVYPIGIEP KEIAKQAAGP LPP-KLAQLK AELKN--VQN IFSVERLDYS KGLPERFLAY EALLEKYPQH HGKIRYTQIA PTSRGDVQAY
AtTPS1         GVEDQGRLTR VAAFPIGIDS DRFIRALEVP EVIQHMKELK ERFAG--RKV MLGVDRLDMI KGIPQKILAF EKFLEENANW RDKVVLLQIA VPTRTDVPEY
SvTPS1         GVEDQGRLTR VAAFPIGIDS DRFKRALELP AVKRHISELT QRFAG--RKV MLGVDRLDMI KGIPQKILAF EKFLEENPAW NDKVVLLQIA VPTRTDVPEY
SvTPS2         GINYHGRTVL VKTFTVGVDM GRLRATLASP EAAAKAKEIA EAYRG--RVL MVGVDDVDPF KGVGLKLLAL EKMLEADQDM RRRVVLVQIN NPARSAGGDA
SvTPS3         GIEYYGRTVT VKILPVGIDM GQLRSVVSAP ETGDVVRRVA EAYKG--RRL MIGVDDVDLF KGIGLKFLAM EQLLVEHPEL RGRAVLVQIA NPARSEGRDV
SvTPS4         GIEYFGRTVN LKILAAGVHV GRLESMLKLP VTISKVQEIE NRYRG--KLV ILGVDDMDIF KGISLKLLGL ELLLERTPKL RGKVVLVQIV NPARSIGKDV
SvTPS5         ALEYYGRTVT IKILPVGVHL EQLQSVLNLP ETGVKVAELL KQFYHRNRLL LLGVDDMDIF KGISLKLLAF EQLLMQHPEW RGRVVLVQIA NPARGKGKDV
SvTPS6         GLDYFGRTVG IKIMPVGINT VQLQSLLQQP DLERQVAELR NQFDR--KTV LLGVDDMDIF KGIDLKILAF EQMLKTHPKW QGRAVLVQIA NPKGGSRKDL
SvTPS7         GLDYFGRTVG IKIMPVGIHM GQLQSGLRLP DREWRLSELQ RQFEG--KTV LLGVDDMDIF KGINLKLLAF ENMLRTHPKW QGRAVLVQIA NPARGKGKDL
SvTPS8         GIEYFGRTVS LKILSVGVHV GRLESVLNLP ATVSKVQEIE QRYKG--KML MLGVDDMDIF KGISLKLLGL ELLLERNPKL RQKVVLVQII NPARSTGKDV
SvTPS9         GIEYYGRTVM VKILPVGIDM GQLRSVVSAP ETGDAVRRVA EAYKG--RRL MIGVDDVDLF KGIGLKFLAM EQLLLEHPEL RGRAVLMQIA NPARSEGRDV
SvTPS10        CLEYYGRTVS IKILPVGVHM EQLKTVLGLP KTEAKVAELM EMYMGKGRVV MLGVDDMDIF KGISLKLLAM EELLRQHPEW RGKLVLVQVA NPARGRGKDV

                       510        520        530        540        550        560        570        580        590        600
               ....|....| ....|....| ....|....| ....|....| ....|....| ....|....| ....|....| ....|....| ....|....| ....|....|
otsA           QDIRHQLENE AGRINGKYGQ LGWTPLYYLN QHFDRKLLMK IFRYSDVGLV TPLRDGMNLV AKEYVAAQDP ---------- -----ANPGV LVLSQFAGAA
AtTPS1         QKLTSQVHEI VGRINGRFGT LTAVPIHHLD RSLDFHALCA LYAVTDVALV TSLRDGMNLV SYEFVACQE- ---------- -----AKKGV LILSEFAGAA
SvTPS1         QKLTSQVHEI VGRINGRFGT LTAVPIHHLD RSLDFHALCA LYAVTDVALV TSLRDGMNLV SYEYVACQG- ---------- -----SKKGV LILSEFAGAA
SvTPS2         DDVRSEAAQI ERRINRRFGD GG-ELVVAID GPVPMWVKAA YYAAADCCVV TAVRDGLNRI PYFYTVCRE- EGA------- -----RRGGA VVVSQFAGCA
SvTPS3         QGVQDEAKAI SARVNARFGT PGYTPIVLID GPLTAHEKAA YYAAAECCVV SAVRDGLNRI PYIYTVCRQ- ESNAL----- -GEDSPKRSV IVLSEFVGCS
SvTPS4         EEAKNEAVSV AQRINDKYGS ANYKPVVLID YSIPFYEKIA FYAASDCCIV NAVRDGMNLI PYEYTVCRQ- GNEDIDKLRG SDKSSLHTST LIVSEFVGCS
SvTPS5         KEVQEESDAM VRRINDAFGQ PDYQPVILID KPLQFYERMA YYVVAEFCLV TAVRDGMNLI PYEYVIARQ- GNEMIDRILG LGPSSRKKSM LVVSEFIGCS
SvTPS6         EELQAEIEVS CKRINEQFGR PGYSPVVLVN RTLSSVERMA YYTIAECVVV TAVRDGMNLT PYEYIVCRQ- GAPGLDGSGD D--RPRGKSM LVVSEFIGCS
SvTPS7         EAIQAEIQES CRRINGDFGQ SGYSPVVFID RDVSSVEKIA YYTIAECVVV TAVRDGMNLT PYEYIVCRQ- GAPGSEAMSE V--SGPKKSM LVVSEFIGCS
SvTPS8         QEAITETVSV AERINRKYGS SGYNPVVLID HHIPFYEKIA FYAASDCCIV NAVRDGMNLV PYEYTVCRQ- GNEEIDKLRG FDKDTSHTST LIVSEFVGCS
SvTPS9         QGVQDEAKAI SARVNERFST AGYTPIVLID GLVTEQEKSA YYAAAECCVV SAVRDGLNRI PYIYTVCRQ- ESNAL----- -GDDSPKRSV IVLSEFVGCS
SvTPS10        AEVQAETYAM VRRINEVYGE PGYEPVVLID EPLQFYERVA YYVIAEVCLV TAVRDGMNLI PYEYIVSRQ- GNEKLDRMLR QGKPEEKKSM LVVSEFIGCS
```

(Continues on next page)

```
                       610        620        630        640        650        660        670        680        690        700
               ....|....| ....|....| ....|....| ....|....| ....|....| ....|....| ....|....| ....|....| ....|....| ....|....|
otsA           NELTS----A LIVNPYDRDE VAAALDRALT MSLAERISRH AEMLDVIVKN DINHWQECFI SDLKQIVP-- ---------- ---------- ----------
AtTPS1         QS---LGAGA ILVNPWNITE VAASIGQALN MTAEEREKRH RHNFHHVKTH TAQEWAETFV SELNDTVIEA QLR------- -----ISKVP PE---LPQHD
SvTPS1         QS---LGAGA ILVNPWNITE VADSIRHALT MSSDEREKRH RHNYAHVTTH TAQDWAETFV CELNDTVAEA QLR------- -----TRQVP PG---LPSQT
SvTPS2         AS---LG-GA IRVNPLDPAA LADAMRAAVT MDAEERQARH SANYGYLRAH DVADWARSFD KALNLACRDH AALLLVGFGL GLGYRAMALR PRFQALTAEH
SvTPS3         PS---LS-GA IRVNPWSVES VAEAMNAALR MPEAEQRLRH EKHYKYVSTH DVAYWARSFD QDLQRACKDH FSRRHWGIGF GMSFKVVALG PNFRRLSVEH
SvTPS4         PS---LS-GA FRVNPWSVED VADALYSATD LTQFEKIQRH EKHYRYVKSH DVTYWARSFD QDLERTCKEQ DSRRCWTTGF GLNFRVIALS PGFRRLSLEH
SvTPS5         PS---LS-GA IRVNPWNIDS VADAMDSALE MPEGEKVLRH EKHHRYVSTH DVGYWANSFL QDLERTCLDH NRRRCWGIGF GLKFRVVALD PNFKKLAVEH
SvTPS6         PS---LS-GA IRVNPWNIES TAEAMNESIA LSDNEKQLRH EKHYRYVSSH DVAYWSKSFI HDFERSCRDH FRRRCWGVGL GFGFRVVALD RNFKKLTVDS
SvTPS7         PS---LS-GA IRVNPWNIEA TAEGMNEAIS MPEQEKHLRH EKHYRYVSSH DVAYWSKSFI QDLERACKDH FRRTCWGIGL GFGFRVVALD PHFTKLNIDS
SvTPS8         PS---LS-GA FRVNPWSVDD VADALCHATD LTESEKRLRH EKHYRYVSTH DVAYWARSFA QDLERACKDH YSRRCWAIGF GLNFRVIALS PGFRKLSSEH
SvTPS9         PS---LS-GA IRVNPWSVES VAEAMNAALQ MPEAEQRLRH EKHYKYVSTH DVAYWARSFD QDLQRASKDH LSRRHWGIGF GMSFKVVALG PNFRRLYVKH
SvTPS10        PS---LS-GA VRVNPWNIEA VADAMESALV LPENEKKLRH DKHFRYVSTH DVGYWANSFL QDLERTCKDH AHRRCWGIGF GLRFRVVSLD RHFKKLSLES

                       710        720        730        740        750        760        770        780        790        800
               ....|....| ....|....| ....|....| ....|....| ....|....| ....|....| ....|....| ....|....| ....|....| ....|....|
otsA           ---------- ---------- ---------- ---------- ---------- ---------- ---------R SAESQQ---- ---------- ----------
AtTPS1         AIQRYSKSNN RLLILGFNAT LTEPVDNQGR RG-DQIKEMD LNLHPELKGP LKALCSDPST TIVVLSGSSR SVLDKNFGE- YDMWLAAENG MFLRLTNG-E
SvTPS1         AIQQYLRSKN RLLILGFNST LTEPVESSGR RGGDQIKEME LKLHPDLKGP LKALCEDEHT TVIVLSGSDR SVLDENFGE- FKMWLAAEHG MFLRPTYG-E
SvTPS2         VRPSYRDAAC RLILLDYDGT MVPEQQDER- ---------K GAPDAGIVRL LNELCADPKN VVFLVSGRGK DQLAAWFASC EKLGICAEHG YFTRWSKGDP
SvTPS3         IVPSYRRTEN RLILLDYDGT VMPENSI--- ---------D KTPSSEVISV LNRLCEDPKN RVFIVSGRGK DELSKWFAPC EKLGIAAEHG YFTRWSRDAP
SvTPS4         FASSYKKANR RVIFLDYDGT LVPQSSL--- ---------N KAPSAEVISI LNSLCNDTKN TVFIVSGRGR NSLSEWFDSC ENLGIAAEHG YFIRWNKAAE
SvTPS5         LVSAYRRTTK RVILLDYDGT LMPQTSF--- ---------G KSPTSKTIDM LNSLCRDKNN MIFLVSTKSR MTLNEWFSPC ENLGLAAEHG YFLRLRRDAE
SvTPS6         IVADYKKSKS RVILLDYDGT LIPQTTM--- ---------N KTPNETVVSM MNTLCADKKN VVFIVSGRGR DSLEKWFYPC PELGIAAEHG YFMRWTRDEQ
SvTPS7         IVNAYEISES RAILLDYDGT LVPQTSI--- ---------N KAPSPEVLSI INTLCSDRRN IVFLVSGRDK DTLGEWFASC PKLGIAAEHG YFLRWSREEE
SvTPS8         FVSCYNKASR RAIFLDYDGT LVPQSSI--- ---------N KAPSAEVISI LKTLCNDPKN NVFIVSGRGR DSLDEWFSPC EKLGIAAEHG YFVRWSKEAE
SvTPS9         IVPSYRKTEN RLILLDYDGT VMPENSI--- ---------D KTPSSEVISV LNCLCEDPKN RVFIVSGRGK DELSKWFAPC EKLGIAAEHG YFTRWSRDTP
SvTPS10        ILIAYRNAKT RAILLDYDGT LMPQA-I--- ---------N KSPSAESVRI LNSLCRDKNN EVYLCSGYDR RTLHEWF-PC ENLGIAAEHG YFLRSKRDAE

                       810        820        830        840        850        860        870        880        890        900
               ....|....| ....|....| ....|....| ....|....| ....|....| ....|....| ....|....| ....|....| ....|....| ....|....|
otsA           ---------- ---------- ---------- -RDKVATFPK LA-------- ---------- ---------- ---------- ---------- ----------
AtTPS1         WMTT---MP- -EHLNMEWVD SVKHVFKYFT ERTPRSHFET RDTSLIWNYK YADIEFGRLQ ARDLLQHLWT GPISNASVDV VQGSRSVEVR AVGVTKGAAI
SvTPS1         WMTT---MP- -EHLNMDWVD SVKHVFEYFT ERTPRSHFEH RETSFVWNYK YADVEFGRLQ ARDMLQHLWT GPISNAAVDV VQGSRSVEVR SVGVTKGAAI
SvTPS2         WEVMNKQVVP G-GTGSGWKA VAERVMRQYM EATAGSYMET KETALAWHYQ NADPVVGPCQ AKELHDHLV- GLLADEPASV AIGSQIVEVN PQGVNKGVAV
SvTPS3         WETS---VL- --AADFDWKK TAEPVMQLYT GATDGSYIEH KESAIVWHHH EADPDFGSCQ AKELLDHLE- NVLANEPVVV KRGQHIVEVN PQGISKGVVV
SvTPS4         WETS---SS- --GQCSEWKL IADPVMHVYT ETTDGSSIEC KESALVWHYQ NTDHDFGSCQ AKELVSHLE- RVLANEPVVV KRGHQIVEVK PQGVSKGIAV
SvTPS5         WETC---VP- --VIDCSWKQ IAELVMKTYT ETTDGSTIED KETAIVWSYE DADPDFGSCQ AKELHDHLE- SVLSNEPVSV KAGLNHVEVK PQGVSKGLVA
SvTPS6         WQIQ---NP- --TSEFGWMH MAEPVMKLYT EATDGSYIET KESALVWHHQ DADPGFGSSQ AKEMLDHLE- SVLANEPVSV KSGQHIVEVK PQAVSKGFVA
SvTPS7         WQTC---TQ- --TLDFGWMQ MAKPVMNLYT EATDGSSIET KESALVWHHQ DADPGFGSSQ AKEMLDHLE- SVLANEPVSV KSGQFIVEVK PQGVSKGVVA
SvTPS8         WESS---YP- --RTQREWKH IAEPVMKVYT ETTDGSSIEP KESALVWHYL DADHDFGSCQ AKELQDHLE- RVLSNEPVVV KCGHYIVEVK PQGVSKGLAV
SvTPS9         WEIS---VL- --AADFDWKK TAEPVMQLYT EATDGSYIEH KESAIVWHHH EADPDFGSCQ AKELLDHLE- NVLANEPVVV KRGQHIVEVN PQGISKGVVA
SvTPS10        WQTC---IT- --PADCSWKQ IAEPVMCLYR ETTDGSTIED RETILVWNYE DADPDFGSCQ AKELVDHLE- SVLANEPVSV RTTPHSVEVK PQGVSKGLVA

                       910        920        930        940        950        960        970        980        990       1000
               ....|....| ....|....| ....|....| ....|....| ....|....| ....|....| ....|....| ....|....| ....|....| ....|....|
otsA           ---------- ---------- ---------- ---------- ---------- ---------- ---------- ---------- ---------- ----------
AtTPS1         DRILGEIVHS KSMTTPIDYV LCIGHFLGKD EDVYTFFEPE LPSDMPAIAR SRPSSDSGAK SSSGDRRPPS KSTHNNNKSG SKSSSSSNSN NNNKSSQRSL
SvTPS1         DRILGEIVHS ENMVTPIDYV LCIGHFLGKD EDIYVFFDPE YPSES----K VKPEGSS--- -TALDRRP-- -----NGRSS NGRSNSRNSQ SRTQKVQQAV
SvTPS2         RGVLSAMSRR G--VAAPDLA LCFGDGEA-D EDMFEALARS G--------- ---------- ---------- ---------- ---------- ----------
SvTPS3         ESLLSSMVKT G--K-PPDFV LCIGDDRS-D EDMFESIVCP S--------- ---------- ---------- ---------- ---------- ----------
SvTPS4         DKIIRTLVSK G--E-VADLL MCIGNDRS-D EDMFESINKA T--------- ---------- ---------- ---------- ---------- ----------
SvTPS5         KRILSTMQER G--D-LPDFI LCVGDDRS-D EDMFEVITAA A--------- ---------- ---------- ---------- ---------- ----------
SvTPS6         EKILSTLMEK G--R-QADFV LCIGDDRS-D EDMFEQISDI M--------- ---------- ---------- ---------- ---------- ----------
SvTPS7         ERILASMKER G--K-QADFV LCIGDDRS-D EDMFENIADI M--------- ---------- ---------- ---------- ---------- ----------
SvTPS8         DKLIRSLVNN G--K-APDFL MCIGNDRS-D EDMFESINGM T--------- ---------- ---------- ---------- ---------- ----------
SvTPS9         DSLLSSMVKT G--K-PPDFV LCIGDDRS-D EDMFESIVCP S--------- ---------- ---------- ---------- ---------- ----------
SvTPS10        RRMLASMQER G--Q-CPDFV LCIGDDKS-D EDMFQLIATA A--------- ---------- ---------- ---------- ---------- ----------

                       1010       1020       1030       1040       1050       1060       1070       1080       1090      1100
               ....|....| ....|....| ....|....| ....|....| ....|....| ....|....| ....|....| ....|....| ....|....| ....|....| ....|
otsA           ---------- ---------- ---------- ---------- ---------- ---------- ---------- ---------- ---------- ---------- -----
AtTPS1         QSERKSGSNH SLGNSRRPSP EKISWNVLDL KGENYFSCAV GRTRTNARYL LGSPDDVVCF LEKLADTTSS P--------- ---------- ---------- -----
SvTPS1         S-ERSSSSSH SSTSSDHSWR E--GSSVLDL KGENYFSCAV GRKRSNARFL LSSSEDVVSF LKELATATAG FQSSNADYMF LDRQ------ ---------- -----
SvTPS2         ---------- ---------- ------ARLA AGARVFACTV GKKATEAAFY VEEPADVVGL LRALARCYSY ---------- ---------- ---------- -----
SvTPS3         ---------- ---------- --NA-SVKLP ATSEVFACTV GKKPSMAKYY LDDTVDVIKM LEGLANAPSQ RPRPAVQLRV SFEGSL---- ---------- -----
SvTPS4         ---------- ---------- --SLA--ELP AIPEVFACSV GPKASKANYY VDGCSEVIRL LKGVIDVSSQ KDTTSHS-HV NSNDILEVVS ---------- -----
SvTPS5         ---------- ---------- --ARGPSPLH PEAEVFACTV GRKPSKAKYY LDDSADIVRL IQGLASVSDD QALHGGA-PL PNAAAATDTI PR-------- -----
SvTPS6         ---------- ---------- --RR--SMVD PQTSLYACTV GQKPSKAIYY LDDANDVLNM LEALADASEE AGSGSPE-AT EEEGPLTLEQ A--------- -----
SvTPS7         ---------- ---------- --KR--NIVA PRTPLFACTV GQKPSKAKFY VDDTFEVVTM LSALADATEP ELETDSA-DE LVASTLLLNI GDEQSECTDR SIGGS
SvTPS8         ---------- ---------- --SNT-VLSP TVPEVFACSV GQKPSKAKYY VDDTTEVIRL LKNVTRSSSQ REDVSHG-RV TFRDVIDFVE ---------- -----
SvTPS9         ---------- ---------- --NA-SVKLP ATSEVFACTV GKKPSMAKYY LDDTVDVIKM LEALANATSQ RPRLAVQLRV SFEGSL---- ---------- -----
SvTPS10        ---------- ---------- --CG--DSLA SKAEVFACTV GRKPSKAKYY LDDAAEVVRL MQGLSYVSEE LALANHR-DE DEDSS-LDVW E--------- -----
```

**Supplementary Figure S4. Multiple sequence alignment of *S. viridis* TPS proteins.** Residues predicted to be involved in UDPGlc or Glc6P binding based on comparison with crystal structures of otsA in complex with UDP and Glc6P or UDPGlc (Gibson *et al*., 2002; 2004) are highlighted in blue and red, respectively. The putative sumoylation sites (Fitchner and Lunn; 2021) are highlighted in magenta. The list of coding genes is presented in Supplementary Table S1. otsA, *Escherichia coli* TPS; At, *Arabidopsis thaliana*; Sv, *Setaria viridis*.

```
                        10         20         30         40         50         60         70         80         90        100
               ....|....| ....|....| ....|....| ....|....| ....|....| ....|....| ....|....| ....|....| ....|....| ....|....|
otsB           ---------- ---------- ---------- ---------- ---------- ---------- ---------- ---------- ---------- ----------
SvTPP1         MDMSTSSPV- ---------- -ITDP----- L-SISPPLLG SLAS-NMMQF SVMS---GGC SSP------- --------SM NVSASRRKIE E-VLVNGLLD
SvTPP2         --MTNQDVVV ----SEMGIA AGAA-LPG-- -PR--P---- --ALLACRGA AA-GAMSHRY LDLAAAA--- ---------- -AR---S--- --ASCTW-AD
SvTPP3         --MAKPSVGV ----SEVGVS AAPA-QSACP CPG--T---- LFPYPPPRGA GIAAAVRRKC LQVELGA--G ---------- -AG---L--- --LGGAWGVE
SvTPP4         MGSYANGS-- ---------- ---------- ---------- TCA----NDV PPSA------ ---------- -------EEM KEPAFPLKTM PLHANGW-LN
SvTPP5         --MTKHAAF- ---------- AAED-VAAAA QPG--R---- RFTSYPP--- -ASC--RTAA AQ-------G MVDP--S--- -TAAGVV--- A-RAGSW-LG
SvTPP6         ---------- ---------- ---------- ---------- ---------- ---------- ---------- ---------- ---------- ----------
SvTPP7         --MTKQAVV- ------VPVT PEAA-VAVPP NSA--P---- LFPYPPPRAA APGAAVRKKY LQMGGGG-AG TVGA----GA AAGGGPG--- R-IG-GW-VE
SvTPP8         MDLKTGLN-- SPVL----AD HLPT-----A LPA--A---- VMTFTTPTSF PSPG----LC L------N-- --------TT KKIPLPGKVE EVRATGW-LD
SvTPP9         --MTKHGVV- ---------- LGPE-DAVVA AAA--R---- HFSFPPPRTG ADSC--RKLA AQ-------- -VDLGA---- ---A-VM--- ----GSW-LD
SvTPP10        ---------- ---------- ---------- ---------- -MK------- -AS------- ---------- ---------- ---------- -----S----

                       110        120        130        140        150        160        170        180        190        200
               ....|....| ....|....| ....|....| ....|....| ....|....| ....|....| ....|....| ....|....| ....|....| ....|....|
otsB           ---------- ---------- ---------- ------MTEP LTETPELSA- ----KYAWFF DLDGTLAEIK PHPDQVVVPD NILQGLQLLA TASDGALALI
SvTPP1         AMKSS-SPRK KHNLAF--GQ GNSLDEDPVY SSWMSKCPSA LTSFKQIVAN AQGKKIAVFL DYDGTLSPIV DDPNKAFMSP VMRAAVRN-- VAKSFPTAIV
SvTPP2         AMRAS-SPTR SRAAA----- -----DVDEF TAWMRKHPSA LGKFEQIASS SKGKKIVMFL DYDGTLSPIV ADPDAAYMSD AMRSAVRD-- VAKYFPTAIV
SvTPP3         SMRAS-SPTH AKAAAA---- LAAGVDDER- AAWMVRHPSA LGKFEQIVAA SEGKRIVMFL DYDGTLSPIV DDPDAAFMSE TMRMAVRS-- VAKHFPTAIV
SvTPP4         DMKIS-SPTA IRV-----NI GNPSAFDPIY RAWTKKYPSA LNAFEKIAAY GKGKKTVLFL DYDGTLSPIV DEPDNAIMSD QMREVVRN-- AALHLPTAII
SvTPP5         ---A--VPRR ---------- -----AEAEH DDWMEKHPSA LAAFESVLAA AKGKQVVMFL DYDGTLSPIV KDPDSAVMTD EMRDAVRG-- VAEHFPTAIV
SvTPP6         ---------- ---------- ---------- ---------- ---------- -------MFM DYDGTLSPIV TDPDMAFMTA EMRTSVRD-- VAKHFPTAIV
SvTPP7         SMRAS-SPTH ARAAAA---- LAAGVDVERH ASWMVEHPSA LSKFDQVVAA SKGKQIVVFL DYDGTLSPIV DDPDAAYMSD TMRRAVRS-- VAKHFPTAIV
SvTPP8         LMMAS-SPTR KRQIKDVVND TQADDLDLQY RNWMVDYPSA LTSFETITDL AGSKRLALFL DYDGTLSPIV DNPANALMSD EMRAAVRH-- VASLFPTAII
SvTPP9         SMKASSPRHR LM-AP----- -LAGAADAEH DDWMERHPSA LERFEALAAA AKGKQVAVFL DYDGTLSPIV EDPDRAVMTD EMREAVRG-- VAARFPTAIV
SvTPP10        -------PRR A--------- -----ADAEH GDWMEKHPSA LTWFEPALAA AKGKQIVMFL DYDGTLSPIV EDPDRAVMSE EMRDAVRR-- VAEHFPTAIV
                                                                                |------|
                                                                                 Motif I

                       210        220        230        240        250        260        270        280        290        300
               ....|....| ....|....| ....|....| ....|....| ....|....| ....|....| ....|....| ....|....| ....|....| ....|....|
otsB           SGRSMVELDA LAKPYRFPLA GVHGAERRDI NGKT------ ---------- ---HIVHLPD AIARDISVQL H--------- ---------- ---TVIAQYP
SvTPP1         SGRSRKKVFE FVKLKELYYA GSHGMDIVTS VAEHNTE--- ---------K CKEANLFQPA CEFLPMIDEV S--------- ---------K SLLEVISGIE
SvTPP2         SGRCRDKVRN FVGLSELYYA GSHGMDIKGP SSNP------ ---------- --ESVLCQPA SEFLPVIDEV Y--------- ---------K LLVEKTKSTP
SvTPP3         SGRCRDKVFE FVKLAELYYA GSHGMDIKGP AKASSRH--- -----AK--A KAKGVLFQPA SEFLPMIEEV H--------- ---------E RLAETTRCIP
SvTPP4         SGRSCDKVFD FVKLAELYYA GSHGMDIMGP VGETGSVTGN RSCTNSSMKQ DKEVKIFQAA SEFIPMIDEV F--------- ---------R LLVEKIRGID
SvTPP5         SGRCRDKVFN FVKLAELYYA GSHGMDIKGP TAQSKHA--- ------K--A KAEAVLCQPA SEFLPVIDEV ---------- --------CR ALTATTAAIP
SvTPP6         TGRCVEKVCS FVGLSELYYA GSHGMDIKGP SSKD------ ---------- -DQTVLLQPA REFLPVIDKA F--------- ---------R ALEEETRATP
SvTPP7         SGRCRDKVFE FVKLAELYYA GSHGMDIKGP VKGSRHT--- ------KA-A KAKGVLFQPA SQFLPMIEQV H--------- ---------E SLVEKTKSIP
SvTPP8         SGRSRDKVFD FVKLNELYYA GSHGMDIMGP VRKAADSNG- VECIRSTDSQ GKEVNLFQPA SEFLPMITEV Y--------- ---------E KLDESVKDIV
SvTPP9         SGRCRDKVFG FVGLEELYYA GSHGMDIRGP TADPNNH--- ------GKDQ EAKSVLCQPA AEFLPVIEEA YA-------- ---------A LVGSVEASIP
SvTPP10        SGRCRDKVFN FVKLTELYYA GSHGMDIEGP AKQSNKH--- ------VQAN AEEAVHYQAG SEFLPIIEEH PAKHDVMHVR TCRRRWQVYR TLTAKMESIA
              |--|
            Motif II

                       310        320        330        340        350        360        370        380        390        400
               ....|....| ....|....| ....|....| ....|....| ....|....| ....|....| ....|....| ....|....| ....|....| ....|....|
otsB           GAELEAKGMA FALHYRQAPQ HEDALMTLAQ R-ITQIWPQM ALQQGKCVVE IKPRGTS-KG EAIAAFMQEA PFIGR---TP VFLGDDLTDE SGFAVVNRLG
SvTPP1         GASIENNKFC VSVHYRNVAE KDWQVVARLV NEVLEAFPRL KVTNGRMVLE VRPVIDWDKG KAVEFLLQSL GLNDSENVIP IYIGDDRTDE DAFKVLRERN
SvTPP2         GAKVENNKFC LSVHFRCVDE KRWNALAEQV KAVIKDYPKL KLTQGRKVLE IRPSIMWDKG KALEFLLESL GFANCSDVLP VYIGDDRTDE DAFKVLRKRG
SvTPP3         GAKVENNKFC VSVHFRCVDE KMWGEVSEAV KGVLREYPKL RLTLGRMVLE VRPTIKWDKG KALEFLLESL GFADCTNVLP VYIGDDRTDE DAFKVLRRRG
SvTPP4         GAKVENNKFC VSVHYRNVNE KDWPLVARCT DDILKAYPRL RLSHGRKVLE VRPVIDWNKG KAVEFLLDSL GLADSGNVLP IYIGDDRTDE DAFKVLREDK
SvTPP5         GARVENNKFC LSVHFRCVQE EKWRALEEQV RSVLKEYPDL RLTKGRKVLE IRPSIKWDKG NALQFLLEAL GFADSKNVFP IYIGDDRTDE DAFKVLRNMG
SvTPP6         GARVEHNKFC LSVHFRCVDE KSWSSLAEQV KAVLRDFPEL KLTEGRKVLE IRPSIMWDKG KAVEFLLKSL GFDDRSDVLP VYIGDDRTDE DAFKVLKKRG
SvTPP7         GAKVENNKFC VSVHFRCVDE KSWSALADTV KSVLKDYPKL KLTQGRMVFE VRPTIKWDKG KALEFLLESL GFADCADVLP VYIGDDRTDE DAFKVLRRRG
SvTPP8         GARMEDNKFC VSVHYRNVAE EDYKKVFQRV TAVLEDYPCL RLTHGRKVFE VRPVIDWNKG KAVEFLLESL GLNESEDVLP IYVGDDRTDE DAFKVLKASN
SvTPP9         GAKVENNKFC LSVHFRCVEE AAWGALFERV RAVLRDYPGL RLTQGRKVLE VRPMIKWDKG KALEFLLDAL GYAERSDVFP VYVGDDRTDE DAFKVLRSRG
SvTPP10        GAKVEHNKYC LSVHFRCVQE EEWKAVEEEV RSVLKEYPDL KLTHGRKVLE IRPSIKWDKG KALEFLLKSL GYAGRSDVFP IYIGDDRTDE DAFKVLRGMG
                                                                             |----------------------------------|
                                                                                          Motif III

                       410        420        430        440        450        460
               ....|....| ....|....| ....|....| ....|....| ....|....| ....|....| ...
otsB           GMS------V KIGTGATQAS WRLAGVPDVW SWLEMITTAL QQKRENNRSD DYESFSRSI- ---
SvTPP1         ---CGYGILV SQVPKDTEAF YSLRDPSEVM GFLNSLVRWR KRSL------ ---------- ---
SvTPP2         ---QGLGILV SKCPKETNAS YSLQDPGEVM DFLLRLVEWK RKSSPPPPPI MIR------P RV-
SvTPP3         ---QGVGILV SKHPKETSAN YSLQEPAEVM EFLLRLVEWK RLSRSRARLM SLQ------- ---
SvTPP4         ---RGFGILV SSVPKESHAV YSLVDPSEVM DFLKRLVKWK EEEALE---- ---------- ---
SvTPP5         ---QGVGILV SKIPKETSAS YSLREPSEVK EFLHKLVKSK QRD------- ---------- ---
SvTPP6         ---HGLGILV SKCPKETDAS YSLQDPIEVM EFLVRLVEWK RLRSPSAA-- --R------P RAP
SvTPP7         ---QGVGILV SKHPKDTSAS YSLQEPAEVM EFLLRLVEWE RLSKARPKW- ---------- ---
SvTPP8         ---RGFGILV SSIPKESDAF YSLRDPAEVM DFLRKLAAWK EQST------ ---------- ---
SvTPP9         ---QGAGILV SRFPKETAAS FSLRDPAEVR DFLRRLVDAN AT-------- ---------- ---
SvTPP10        ---QGIGILV SKFPKETAAS YSLRDPAEVK EFLRKMVKGK GDGGPVKMMN DIV------H VDH
```

**Supplementary Figure S5. Multiple sequence alignment of *S. viridis* TPP proteins.** Conserved substrate-binding and active site motifs are highlighted in grey. Motif I, DXDX(T/V)(L/V/I); motif II, (S/T)(GX) in a hydrophobic context; motif III, K(X)16-30(G/S)(D/S)XXX(D/N). The list of coding genes is presented in Supplementary Table S1. otsB, *Escherichia coli* TPP; Sv, *Setaria viridis*.

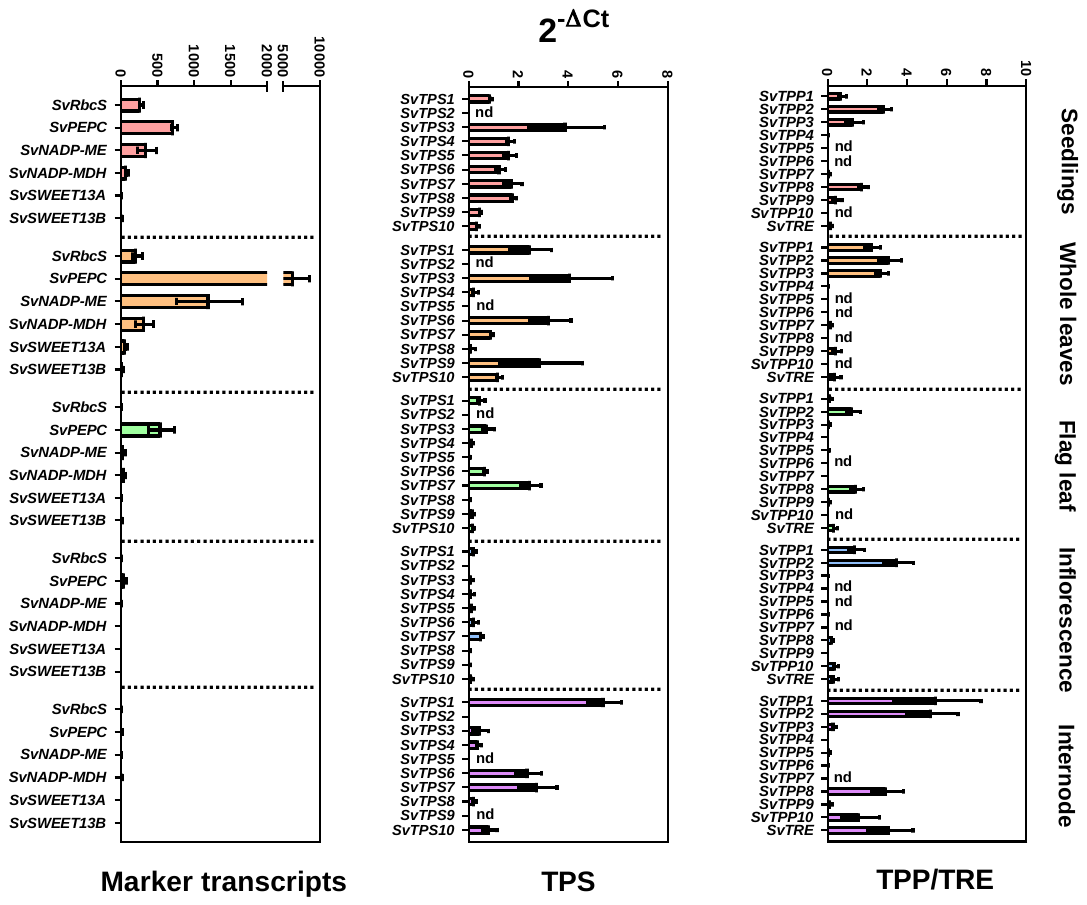

**Supplementary Figure S6. Relative abundance of transcripts encoding Tre6P-related enzymes in different *S. viridis* tissues**. Relative transcript abundance data was calculated with the 2^-ΔCt^ method, using the *SvKIN* transcript as a reference. Data are the mean ± standard error of at least three biological replicates. nd, not detected. The list of coding genes and primers used is presented in Supplementary Table S2.

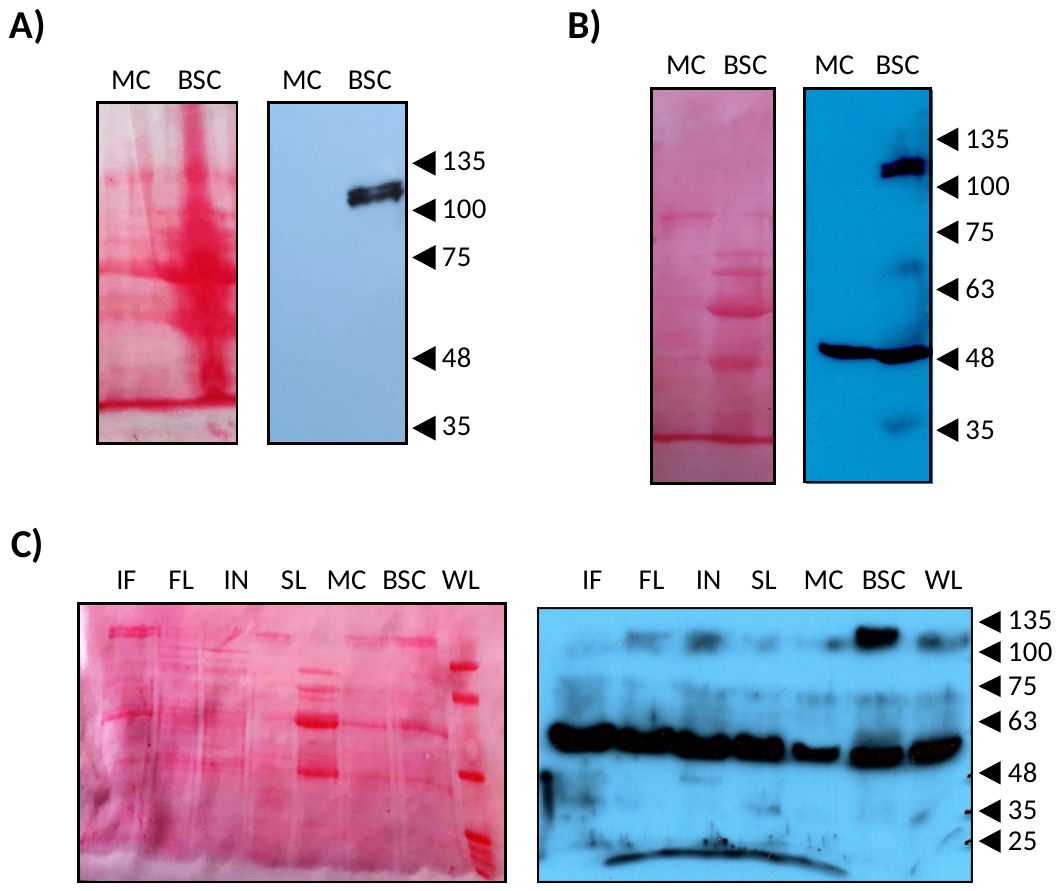

**Supplementary Figure S7. Immunoblotting of SvTPS1 in different tissues and cell types of *S. viridis*.** (A) Immunoblotting of SvTPS1 in MC and BSC from *S. viridis* leaves extracted with TCA. Each lane was loaded with extract from 2 mg fresh weight of tissue. (B) Immunoblotting of SvTPS1 in MC and BSC from *S. viridis* leaves extracted with Laemmli buffer. Each lane was loaded with extract from 2 mg fresh weight of tissue. (C) Immunoblotting of SvTPS1 in different tissues and cell types from *S. viridis*. Each lane was loaded with 25 µg of protein. IF; inflorescence, FL; flag leaf, IN; internode, SL; seedlings, MC; mesophyll cells, BSC; bundle sheath cells, WL; whole leaf. Left panels, membranes stained with Ponceau Red; right panels, X-ray films revealed with ECL. This figure supports Fig. 3 of the main text.

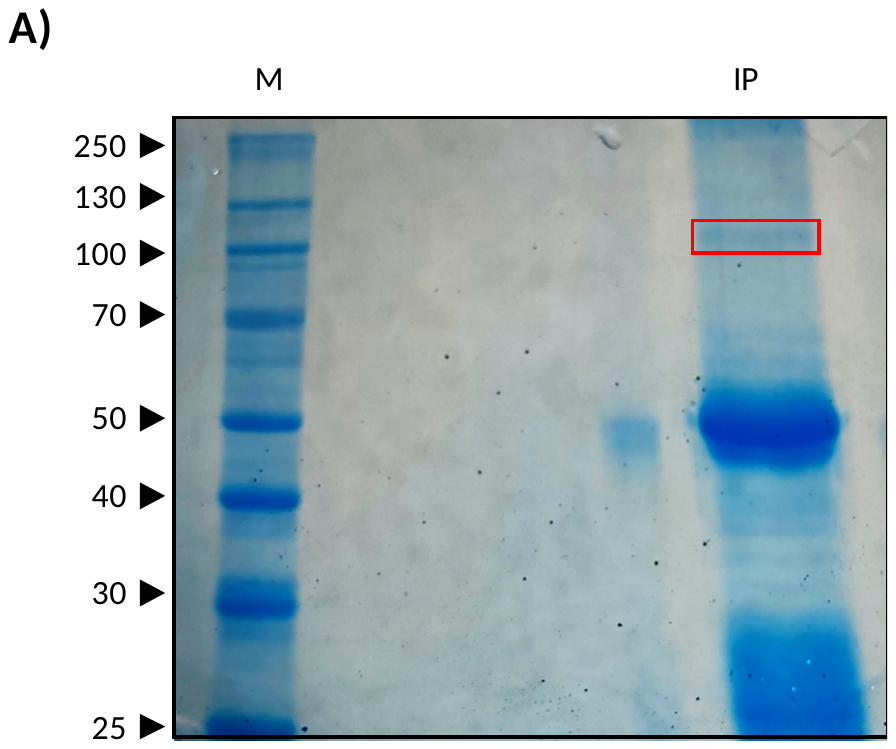

**B)**

```
                 10        20        30        40        50        60        70        80        90       100
        ....|....|....|....|....|....|....|....|....|....|....|....|....|....|....|....|....|....|....|....|
SvTPS1  MSSDAAGGQRSSRNRGRADAAPMPTSSPFTGDGGGAGSPTRVERMLREREHSRRHIFASDTMDTDAAEPVFASAGAFAADGVQSPGRASPANMEDAGGAA

                110       120       130       140       150       160       170       180       190       200
        ....|....|....|....|....|....|....|....|....|....|....|....|....|....|....|....|....|....|....|....|
SvTPS1  SGHAARPPLAGSRSGFRRLGLRGMKQRLLVVANRLPVSANRRGEDQWSLEISAGGLVSALLGVKDVDAKWIGWAGVNVPDEVGQRALTRALAEKRCIPVF

                210       220       230       240       250       260       270       280       290       300
        ....|....|....|....|....|....|....|....|....|....|....|....|....|....|....|....|....|....|....|....|
SvTPS1  LDEEIVHQYYNGYCNNILWPLFHYLGLPQEDRLATTRNFESQFDAYKRANQMFADVVYQHYQEGDVIWCHDYHLMFLPKCLKDHDNNMKVGWFLHTPFPS

                310       320       330       340       350       360       370       380       390       400
        ....|....|....|....|....|....|....|....|....|....|....|....|....|....|....|....|....|....|....|....|
SvTPS1  SEIYRTLPSRLELLRSVLCADLVGFHTYDYARHFVSACTRILGLEGTPEGVEDQGRLTRVAAFPIGIDSDRFKRALELPAVKRHISELTQRFAGRKVMLG

                410       420       430       440       450       460       470       480       490       500
        ....|....|....|....|....|....|....|....|....|....|....|....|....|....|....|....|....|....|....|....|
SvTPS1  VDRLDMIKGIPQKILAFEKFLEENPAWNDKVVLLQIAVPTRTDVPEYQKLTSQVHEIVGRINGRFGTLTAVPIHHLDRSLDFHALCALYAVTDVALVTSL

                510       520       530       540       550       560       570       580       590       600
        ....|....|....|....|....|....|....|....|....|....|....|....|....|....|....|....|....|....|....|....|
SvTPS1  RDGMNLVSYEYVACQGSKKGVLILSEFAGAAQSLGAGAILVNPWNITEVADSIRHALTMSSDEREKRHRHNYAHVTTHTAQDWAETFVCELNDTVAEAQL

                610       620       630       640       650       660       670       680       690       700
        ....|....|....|....|....|....|....|....|....|....|....|....|....|....|....|....|....|....|....|....|
SvTPS1  RTRQVPPGLPSQTAIQQYLRSKNRLLILGFNSTLTEPVESSGRRGGDQIKEMELKLHPDLKGPLKALCEDEHTTVIVLSGSDRSVLDENFGEFKMWLAAE

                710       720       730       740       750       760       770       780       790       800
        ....|....|....|....|....|....|....|....|....|....|....|....|....|....|....|....|....|....|....|....|
SvTPS1  HGMFLRPTYGEWMTTMPEHLNMDWVDSVKHVFEYFTERTPRSHFEHRETSFVWNYKYADVEFGRLQARDMLQHLWTGPISNAAVDVVQGSRSVEVRSVGV

                810       820       830       840       850       860       870       880       890       900
        ....|....|....|....|....|....|....|....|....|....|....|....|....|....|....|....|....|....|....|....|
SvTPS1  TKGAAIDRILGEIVHSENMVTPIDYVLCIGHFLGKDEDIYVFFDPEYPSESKVKPEGSSTALDRRPNGRSSNGRSNSRNSQSRTQKVQQAVSERSSSSSH

                910       920       930       940       950       960       970
        ....|....|....|....|....|....|....|....|....|....|....|....|....|....|..
SvTPS1  SSTSSDHSWREGSSVLDLKGENYFSCAVGRKRSNARFLLSSSEDVVSFLKELATATAGFQSSNADYMFLDRQ
```

**Supplementary Figure S8. Analysis by mass spectrometry of immunoprecipitated SvTPS1.** (A) Immunoprecipitation of SvTPS1 from *S. viridis* BSC analysed by SDS-PAGE. The band highlighted with the red box was sliced from the gel and analysed by LC-MS. (B) Experimentally detected peptides (highlighted in red) were matched to the SvTPS1 protein sequence (UniProt ID A0A4U6VAG9). Coverage: 36%; number of peptides: 28; number of spectra: 51. Data for each peptide are available in Supplementary Table S6.

**Supplementary Figure S9. Immunolocalization of SvTPS1 in cross-sections of *S. viridis* leaves.** Top row, anti-SvTPS1 (1/100); middle row, anti-RbcL (1/1000); bottom row, negative control (without primary antibody). The secondary antibody (anti-rabbit IgG, 1/200) was conjugated with Cy2. The green channel shows the Cy2 fluorescence and the merged images show the Cy2 signal superimposed on the bright-field images. Bars=50 µm. The panel shows representative images of three independent experiments (n=3). bs, bundle sheath; vb, vascular bundle; m, mesophyll; ue, upper epidermis; le, lower epidermis.

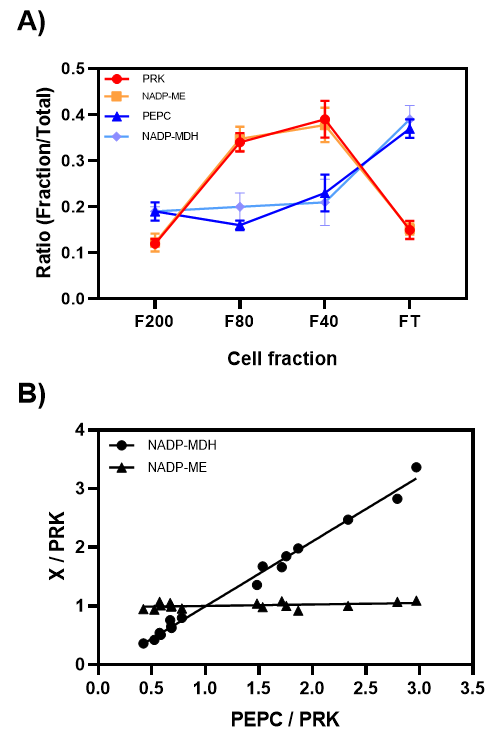

**Supplementary Figure S10. Activities of marker enzymes in different fractions from *S. viridis* leaves.** (A) Activities of PRK, NADP-ME, PEPC and NADP-MDH were measured in fractions of *S. viridis* leaves obtained by sequential sieving of frozen tissue powder through nylon meshes of decreasing pore size. The F200, F80, and F40 fractions correspond to the samples retained on the 200, 80, and 41 μm meshes, respectively, and the FT (flow through) fraction corresponds to the material that passed through all of the meshes. The ratios were calculated by dividing the activity in each fraction by the total activity (i.e. the sum of all the fractions). Data are the mean ± standard error of 4 biological replicates from Experiment 1 (Supplementary Table S3). (B) To test the separation efficiency, regression plots were performed for the marker enzymes: NADP-MDH/PRK or NADP-ME/PRK *vs* PEPC/PRK. In this case, each point corresponds to one biological replicate (Experiment 1, Supplementary Table S3).

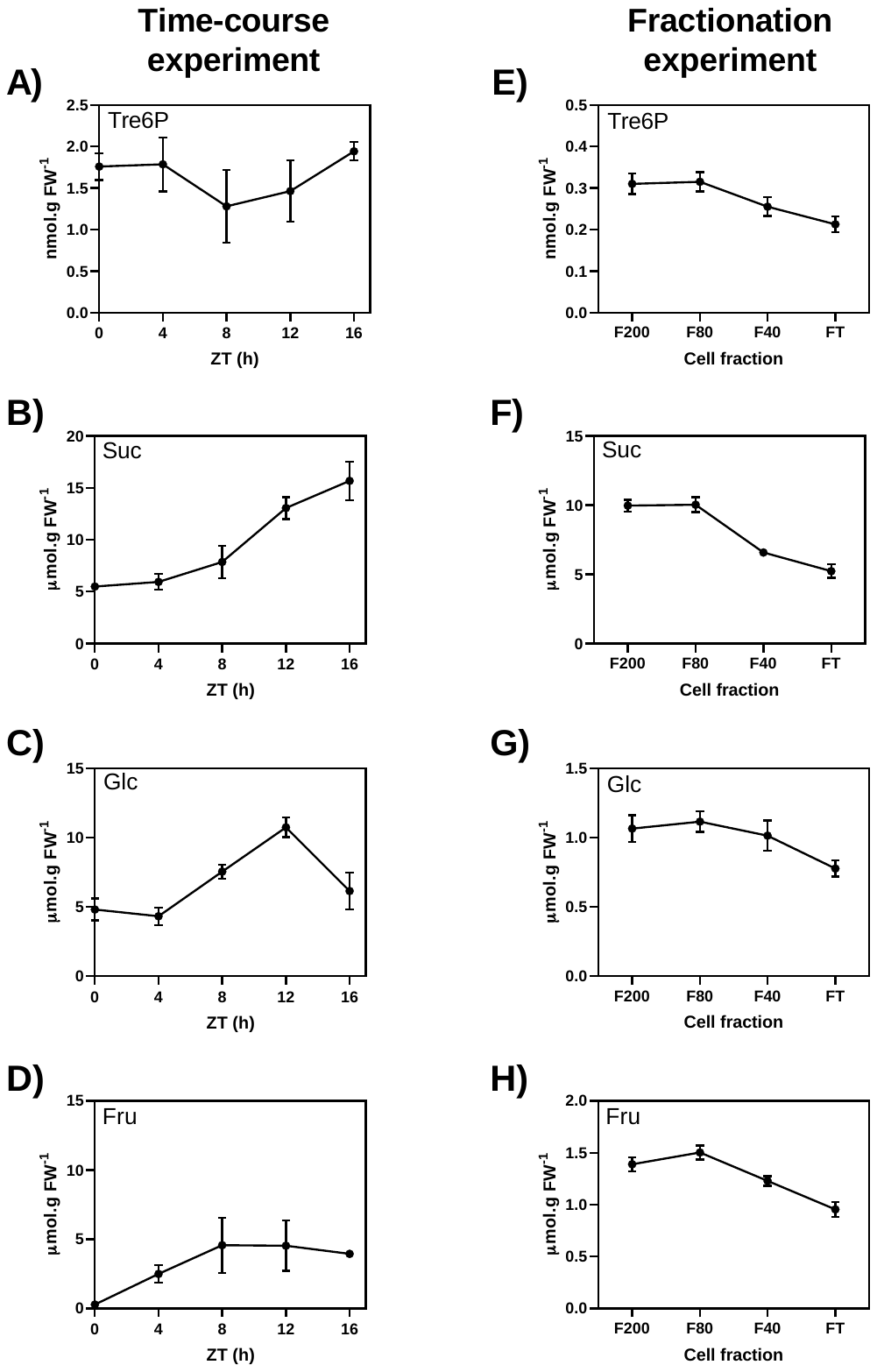

**Supplementary Figure S11. Analysis of metabolites from *S. viridis* leaves.** Data presented in panels A-D correspond to whole-leaf samples from the time-course experiment (Supplementary Table S7), while data presented in panels E-H correspond to one representative fractionation experiment (Experiment 2, Supplementary Table S4). Data are the mean ± SE of 4 biological replicates (n=4).

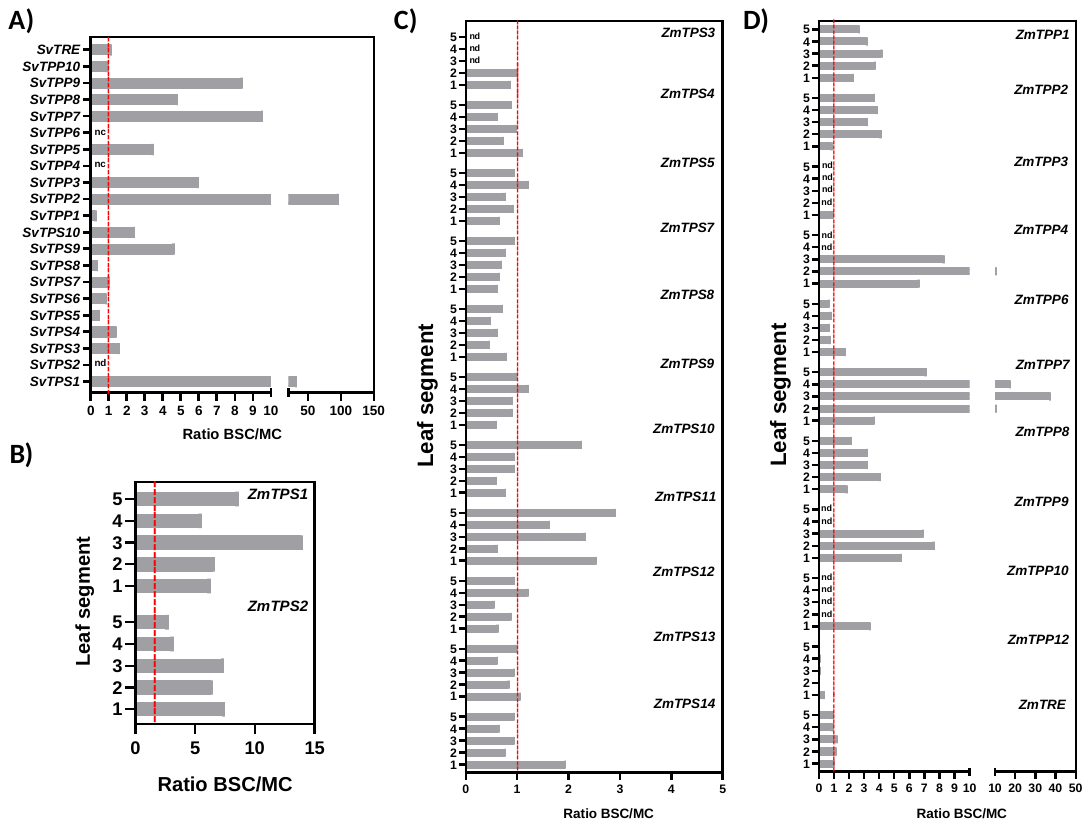

**Supplementary Figure S12. Analysis of transcripts encoding Tre6P-related enzymes in BSC and MC from *S. viridis* and maize leaves.** RNA-seq data of transcripts per million (TPM) from *S. viridis* (A) and maize (B) were retrieved from Supplemental Dataset S3 from Denton *et. al.* (2017), which includes data from John *et. al.* (2014). The ratios were calculated as TPM in BSC/TPM in MC (see Supplementary Table S8 for details). The maize data are derived from *Zea mays* B73 leaf sections representing five contiguous 4-cm slices from tissue just emerging from the ligule (slice 5) to the leaf tip (slice 1). nd, the transcript was neither detected in MC nor BSC; nc, not calculated because the transcript was not detected in MC or BSC.

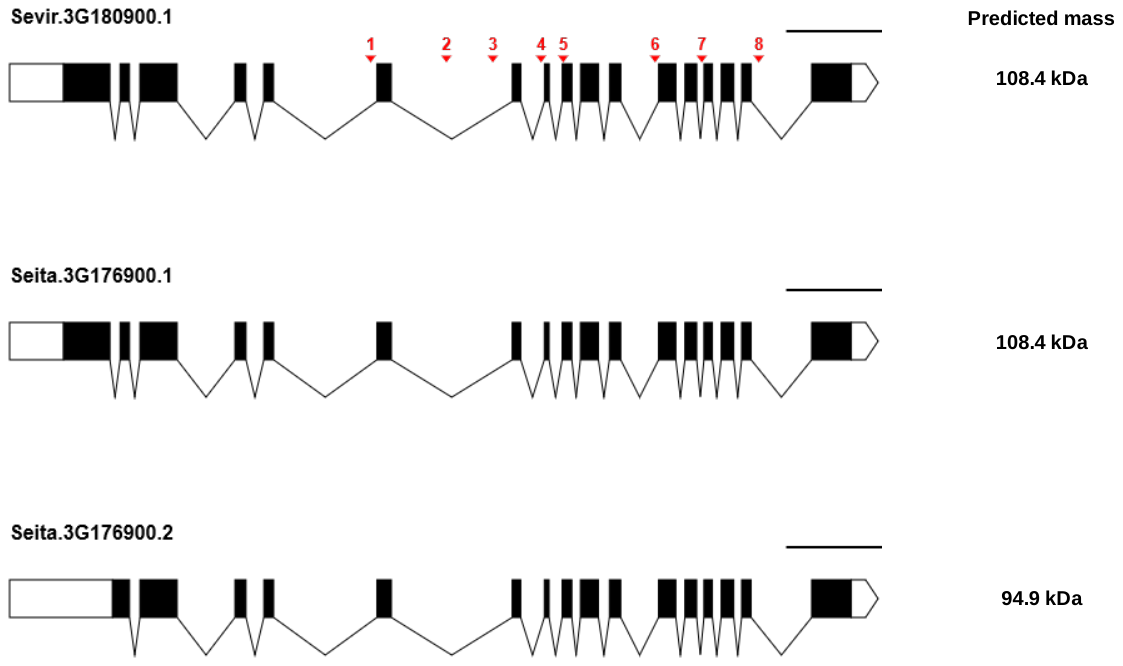

**Supplementary Figure S13. Gene models for the class I TPS from *S. viridis* and *S. italica*.** DNA sequences encoding the class I TPS from *S. viridis* (*Sevir.3G180900.1*, genome version 2.1, Phytozome genome ID 500) and *S. italica* (*Seita.3G176900.1* and *Seita.3G176900.2*, genome version 2.2, Phytozome genome ID 312) were obtained from Phytozome 13 (https://phytozome-next.jgi.doe.gov/). Figures were prepared using the Exon-Intron Graphic Maker tool from WormWeb.org (http://wormweb.org/exonintron). Gene sequences are shown from the transcription start site (left) to the transcription termination site (right), with exons represented by boxes (white = 5´- and 3´- untranslated regions; black = protein-coding regions) and introns represented by lines. Differences between *Sevir.3G180900.1* and *Seita.3G176900.1* are labelled from 1 to 8 and highlighted in red. Scale bars: 1000 bp. The resulting protein sequences were used to calculate the theoretical molecular masses (values presented on the right).
