## Supplementary File S1 for "Intercellular compartmentation of trehalose 6-phosphate metabolism in *Setaria viridis* leaves"

**Supplementary File S1.** Optimised DNA sequence for the expression of SvTPS1 in *E. coli* cells.

> Sevir.3G180900

catATGTCTTCTGATGCAGCAGGCGGCCAGCGTAGCTCTCGTAACCGTGGTCGTGCGGATGCAGCGCCGATGCCGACCAGCTCTCCGTTCACCGGTGATGGTGGTGGTGCTGGCTCTCCGACCCGTGTTGAACGTATGCTGCGTGAACGTGAACACAGCCGTCGTCACATCTTCGCGTCTGATACCATGGACACTGATGCTGCTGAACCGGTTTTCGCGTCTGCTGGTGCTTTCGCGGCTGACGGTGTTCAGTCACCGGGTCGTGCGTCTCCGGCTAACATGGAAGACGCGGGTGGCGCAGCCTCTGGCCACGCAGCGCGTCCGCCGCTGGCTGGTAGCCGTAGCGGCTTCCGTCGCCTGGGTCTGCGCGGCATGAAACAGCGTCTGCTCGTTGTGGCGAACCGTCTGCCGGTCTCTGCCAACCGCCGTGGTGAAGATCAGTGGTCCCTGGAAATCTCCGCTGGCGGTCTGGTGTCTGCTCTGCTGGGCGTGAAAGACGTTGACGCGAAATGGATTGGCTGGGCAGGCGTTAACGTTCCGGATGAAGTAGGTCAGCGTGCACTGACCCGTGCCCTGGCCGAAAAACGTTGCATCCCGGTGTTCCTGGATGAGGAAATCGTTCATCAGTATTACAACGGTTACTGTAACAACATCCTGTGGCCGCTGTTCCACTACCTGGGCCTGCCGCAGGAAGACCGTCTGGCGACCACCCGTAACTTCGAATCCCAGTTCGATGCGTACAAACGTGCGAACCAGATGTTTGCAGACGTAGTCTACCAGCACTACCAGGAAGGCGACGTGATCTGGTGCCACGATTACCACCTGATGTTCCTGCCGAAATGCCTGAAAGACCATGATAACAACATGAAAGTAGGCTGGTTCCTGCACACCCCGTTCCCGTCCTCTGAGATCTATCGTACGCTGCCGTCGCGTCTGGAATTGCTTCGTTCTGTTCTGTGTGCGGACCTGGTAGGCTTCCACACTTACGACTACGCGCGTCACTTCGTTTCCGCGTGCACCCGTATCCTGGGCCTGGAAGGCACTCCGGAAGGTGTTGAAGACCAAGGTCGTCTGACGCGTGTTGCCGCCTTTCCGATCGGTATTGACTCTGATCGCTTCAAACGCGCTCTGGAACTCCCGGCAGTTAAACGTCACATCTCCGAATTAACTCAGCGCTTCGCTGGTCGTAAAGTGATGCTGGGTGTGGACCGTCTGGACATGATCAAAGGTATCCCTCAGAAAATTCTGGCCTTTGAAAAGTTCCTGGAGGAGAACCCCGCGTGGAACGACAAAGTTGTTCTGCTGCAGATTGCGGTACCGACTCGTACTGACGTTCCGGAATACCAGAAACTGACTTCTCAGGTTCACGAAATCGTTGGTCGTATCAATGGCCGTTTCGGTACGCTGACGGCTGTTCCAATTCATCACCTGGACCGTAGCCTTGACTTCCATGCTCTGTGCGCACTGTATGCGGTTACCGACGTGGCGCTGGTTACTAGCCTGCGTGATGGTATGAACCTGGTTTCTTACGAATATGTCGCGTGCCAGGGTTCTAAAAAAGGTGTTCTGATCCTCTCCGAATTTGCGGGTGCAGCGCAATCCCTGGGCGCAGGCGCTATCCTGGTAAACCCGTGGAACATCACTGAAGTCGCGGACTCCATTCGTCACGCGCTGACTATGTCTTCTGATGAACGCGAAAAACGTCACCGCCATAACTATGCACACGTTACCACCCATACCGCGCAGGATTGGGCAGAGACCTTCGTTTGCGAACTGAACGACACCGTGGCTGAAGCCCAGCTGCGCACGCGTCAGGTACCGCCGGGTCTGCCGAGCCAGACCGCTATCCAGCAGTACCTGCGTTCCAAAAACCGCCTGTTGATCCTGGGCTTTAACTCCACGCTGACTGAACCGGTTGAAAGCAGCGGTCGTCGTGGTGGTGATCAGATTAAAGAGATGGAACTGAAACTGCACCCGGATCTGAAAGGTCCGCTTAAGGCGCTGTGCGAAGACGAACACACCACCGTTATCGTTCTGTCCGGTTCCGATCGTTCCGTACTGGATGAAAACTTTGGCGAATTCAAGATGTGGCTGGCAGCTGAGCATGGTATGTTCCTGCGTCCTACCTATGGTGAATGGATGACAACCATGCCGGAACACCTGAACATGGACTGGGTGGATTCTGTTAAACACGTTTTCGAATATTTCACTGAACGTACCCCGCGTTCCCACTTCGAACACCGCGAAACTTCCTTCGTGTGGAACTACAAATATGCTGATGTTGAATTCGGTCGTCTGCAGGCTCGTGACATGCTGCAGCACCTGTGGACCGGTCCGATTAGCAACGCGGCTGTGGACGTTGTTCAGGGTTCTCGTTCCGTTGAAGTTCGTTCCGTTGGCGTCACGAAAGGTGCGGCTATTGATCGCATCCTGGGCGAAATCGTACACAGCGAAAACATGGTTACCCCGATCGATTACGTTCTGTGCATTGGTCACTTCCTGGGTAAAGACGAAGATATCTACGTGTTCTTCGATCCTGAGTATCCGAGCGAATCTAAAGTTAAGCCGGAAGGTTCTTCCACCGCGTTGGACCGTCGCCCGAACGGCCGTTCCTCTAACGGTCGTTCCAACTCCCGTAACTCTCAATCTCGTACGCAGAAAGTTCAGCAAGCGGTGAGCGAACGTTCCTCTTCTTCCTCCCACTCCTCTACTTCCAGCGATCATTCTTGGCGTGAAGGTTCTTCTGTTCTGGACCTGAAAGGTGAAAACTACTTCAGCTGCGCAGTAGGCCGTAAACGTTCTAACGCGCGTTTCCTGCTGTCTTCTTCCGAAGACGTAGTGTCTTTCCTGAAAGAATTAGCGACCGCGACCGCAGGCTTCCAGTCTTCTAACGCTGACTACATGTTCCTGGATCGTCAGctcgagtaagagctc
